# Metabolic effects of the schizophrenia-associated 3q29 deletion are sex-specific and uncoupled from behavioral phenotypes

**DOI:** 10.1101/2020.09.18.303412

**Authors:** Rebecca M Pollak, Ryan H Purcell, Timothy P Rutkowski, Tamika Malone, Kimberly J Pachura, Gary J Bassell, Michael P Epstein, Paul A Dawson, Matthew R Smith, Dean P Jones, Michael E Zwick, the Emory 3q29 Project, Stephen T Warren, Tamara Caspary, David Weinshenker, Jennifer G Mulle

## Abstract

The 1.6 Mb 3q29 deletion is associated with developmental and psychiatric phenotypes. Reduced birthweight and a high prevalence of feeding disorders in patients suggest underlying metabolic dysregulation. We investigated 3q29 deletion-induced metabolic changes using the B6.Del16^+/*Bdh1-Tfrc*^ mouse model. We found that B6.Del16^+/*Bdh1-Tfrc*^ animals preferentially use dietary lipids as an energy source. Untargeted metabolomics showed a strong sex-dependent effect of the 3q29 deletion on fat metabolism. A high-fat diet (HFD) partially rescued the 3q29 deletion-associated weight deficit in females, but not males. Untargeted metabolomics after HFD revealed persistent fat metabolism alterations in females. The HFD did not affect B6.Del16^+/*Bdh1-Tfrc*^ behavioral phenotypes, suggesting that 3q29 deletion-associated metabolic and behavioral outcomes are uncoupled. Our data indicate a HFD intervention in 3q29 deletion syndrome may improve weight phenotypes without exacerbating behavioral manifestations. Our study also highlights the importance of assessing sex in metabolic studies and suggests mechanisms underlying 3q29 deletion-associated metabolic phenotypes are sex-specific.

## INTRODUCTION

3q29 deletion syndrome (3q29del) is a rare (∼1 in 30,000) (Kendall et al., 2017; Stefansson et al., 2014) genomic disorder characterized by a typically *de novo* 1.6 Mb deletion on chromosome 3 (hg19, chr3:195725000-197350000) (Ballif et al., 2008; Glassford et al., 2016; Willatt et al., 2005). The 3q29 deletion is associated with neurodevelopmental and neuropsychiatric phenotypes, including mild to moderate intellectual disability (ID) (Ballif et al., 2008; Cox and Butler, 2015; Willatt et al., 2005), a 19-fold increased risk for autism spectrum disorder (ASD) (Itsara et al., 2009; Pollak et al., 2019; Sanders et al., 2015), and a 20-40-fold increased risk for schizophrenia (SZ) (Kirov et al., 2012; Marshall et al., 2017; Mulle, 2015; Mulle et al., 2010; Szatkiewicz et al., 2014). Two independently generated mouse models of the 3q29 deletion show behavioral manifestations, including social interaction and cognitive deficits, and altered acoustic startle reflex, sensorimotor gating, and amphetamine-induced locomotion (Baba et al., 2019; Rutkowski et al., 2019). The range of neurodevelopmental and neuropsychiatric manifestations in 3q29del is consistent with that observed in other copy number variant (CNV) disorders, including 22q11.2 deletion syndrome (McDonald-McGinn et al., 2015; Schneider et al., 2014), 16p11.2 deletion and duplication syndromes (D’Angelo et al., 2016; Hanson et al., 2015), 7q11.23 duplication syndrome (Mervis et al., 2015), and 1q21.1 deletion syndrome (Brunetti-Pierri et al., 2008). These data demonstrate that risk for multiple neuropsychiatric phenotypes is a feature common to many genomic disorders.

There is growing evidence that metabolic alterations can contribute to neurodevelopmental and neuropsychiatric diseases. Many inborn errors of metabolism have neurodevelopmental and neuropsychiatric manifestations, including Wilson’s disease, cerebrotendinous xanthomatosis, Niemann-Pick disease type C, phenylketonuria, and classical galactosemia (Akil and Brewer, 1995; Bilder et al., 2016; Cox, 1999; Doyle et al., 2010; Imrie et al., 2007; Imrie et al., 2002; Korner et al., 2019; Kuiper et al., 2019; Moghadasian, 2004; Patterson et al., 2012; Vanier, 2010; Wraith et al., 2009). Additional molecular studies of the 22q11.2 deletion have highlighted the role of oxidative stress and mitochondrial dysfunction as major contributors to synaptic phenotypes (Fernandez et al., 2019; Gokhale et al., 2019; Napoli et al., 2015). Mitochondrial function, oxidative stress, and small molecule dysregulation have also been implicated in the pathogenesis of idiopathic ASD, bipolar disorder, major depression, and SZ (Andreazza et al., 2013; Ben-Shachar, 2002; Ben-Shachar and Karry, 2008; Ben-Shachar and Laifenfeld, 2004; Ben-Shachar et al., 1995; Bergman and Ben-Shachar, 2016; Brennand et al., 2015; Frye et al., 2016; Frye et al., 2013; Jones and Sies, 2015; Karry et al., 2004; Marazziti et al., 2012; Ming et al., 2012; Morris and Berk, 2015; Norkett et al., 2016; Park et al., 2016; Park and Park, 2012; Pinero-Martos et al., 2016; Rajasekaran et al., 2015; Robicsek et al., 2013; Rosenfeld et al., 2011; Tarlungeanu et al., 2016; Yap et al., 2010), highlighting some etiological similarities between syndromic and idiopathic cases of neuropsychiatric disorders. In light of this evidence linking metabolic function and neuropsychiatric outcomes, it is noteworthy that four of the 21 genes in the 3q29 deletion interval (the SUMO-specific protease *Senp5*, the fatty acid catabolism protein *Bdh1*, the transferrin receptor *Tfrc*, and the phosphatidylcholine biosynthesis protein *Pcyt1a*) are directly involved in metabolism. Individuals with 3q29del report significantly reduced birthweight and a high proportion of feeding problems and failure to thrive compared to the general population (Glassford et al., 2016), and robust weight deficits have been observed in both mouse models of the 3q29 deletion (Baba et al., 2019; Rutkowski et al., 2019). Additionally, our team previously identified sex differences in the degree of the 3q29 deletion-associated weight deficit in B6.Del16^+/*Bdh1-Tfrc*^ mice, with female animals more substantially affected than males (Rutkowski et al., 2019). These data inspired the hypotheses that metabolic disruption may contribute to 3q29 deletion syndrome phenotypes and that there may be sex-specific effects of the 3q29 deletion on metabolic phenotypes.

Although there are established links between recurrent CNV disorders, metabolism, and neurodevelopmental and neuropsychiatric liability, metabolic function has not been interrogated in the context of the 3q29 deletion. Existing evidence of birthweight deficits in humans with 3q29del, weight deficits in 3q29 deletion mouse models, and the metabolic genes contained in the 3q29 interval motivated our investigation of a possible unidentified metabolic disturbance associated with the 3q29 deletion (Figure 1). Furthermore, to determine whether metabolic disruption and adverse behavioral outcomes arise from a shared mechanism, we interrogated the relationship between metabolic function and behavioral phenotypes in our mouse model. Finally, we have specifically considered sex as a modifier of 3q29 deletion metabolic biology, based on our previously reported finding of sex differences in the severity of the 3q29 deletion-associated weight deficit in B6.Del16^+/*Bdh1-Tfrc*^ mice (Rutkowski et al., 2019) (Figure 1). The new results described here have important implications for our mechanistic understanding of phenotype development in 3q29del and help to further elucidate the relationship between metabolism and neurodevelopmental and neuropsychiatric disease risk.

**Figure 1.**
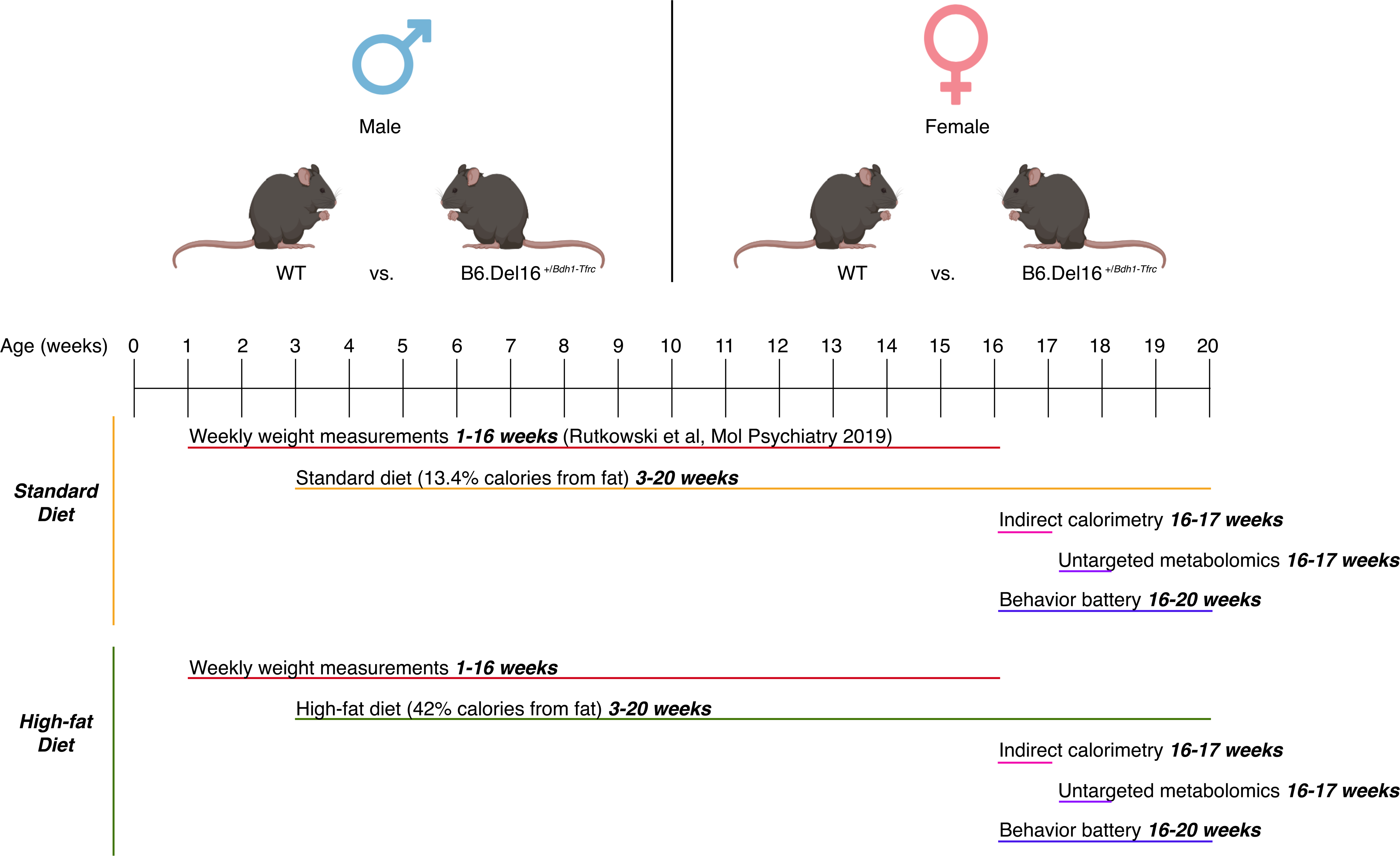
Experimental approach to interrogating the effect of the B6.Del16^+/*Bdh1-Tfrc*^ genotype on metabolism and the effect of sex on 3q29 deletion-associated metabolic phenotypes. All experiments were performed on male and female animals, and the sexes were analyzed separately. Animals were fed either standard diet chow (STD) or high-fat diet chow (HFD) from week3 to week 20. Animals were weighed weekly from week 1 to week 16; STD weights were previously published by our group (Rutkowski et al., 2019). At week 16, a subset of STD- and HFD-treated animals was subjected to indirect calorimetry to assess feeding behavior and metabolic function. At the conclusion of indirect calorimetry, liver tissue was collected for untargeted metabolomics analysis. From weeks 16-20, another subset of STD- and HFD-treated animals was subjected to a behavioral battery.

## RESULTS

### Reduced respiratory exchange ratio and energy expenditure, but not reduced energy consumption, in B6.Del16^+/*Bdh1-Tfrc*^ mice

To investigate whether the 3q29 deletion-associated weight deficit is attributable to increased energy expenditure or decreased energy consumption, we performed 5 days of indirect calorimetry on male and female B6.Del16^+/*Bdh1-Tfrc*^ mice and WT littermates using CLAMS/Metabolic Cages (Columbus Instruments). Energy expenditure was similar between WT and B6.Del16^+/*Bdh1-Tfrc*^ males (Figure 2A); in females, energy expenditure was *reduced* in B6.Del16^+/*Bdh1-Tfrc*^ animals relative to WT (Figure 2B). These data show that the 3q29 deletion-associated weight deficit is not due to increased energy expenditure; rather, B6.Del16^+/*Bdh1-Tfrc*^ mice burn *fewer* calories than their WT littermates. To understand whether the 3q29 deletion influences the calorie source that B6.Del16^+/*Bdh1-Tfrc*^ mice use, we evaluated the respiratory exchange ratio (RER). If an animal is predominantly using lipids as an energy source, the RER will approach 0.7, whereas if an animal is predominantly utilizing carbohydrates, the RER will approach 1 (Meeng et al., 2010). RER was reduced in both male and female B6.Del16^+/*Bdh1-Tfrc*^ mice relative to WT (Figure 2C-D), indicating that B6.Del16^+/*Bdh1-Tfrc*^ animals preferentially use dietary lipids as an energy source, whereas WT animals use more carbohydrates.

**Figure 2.**
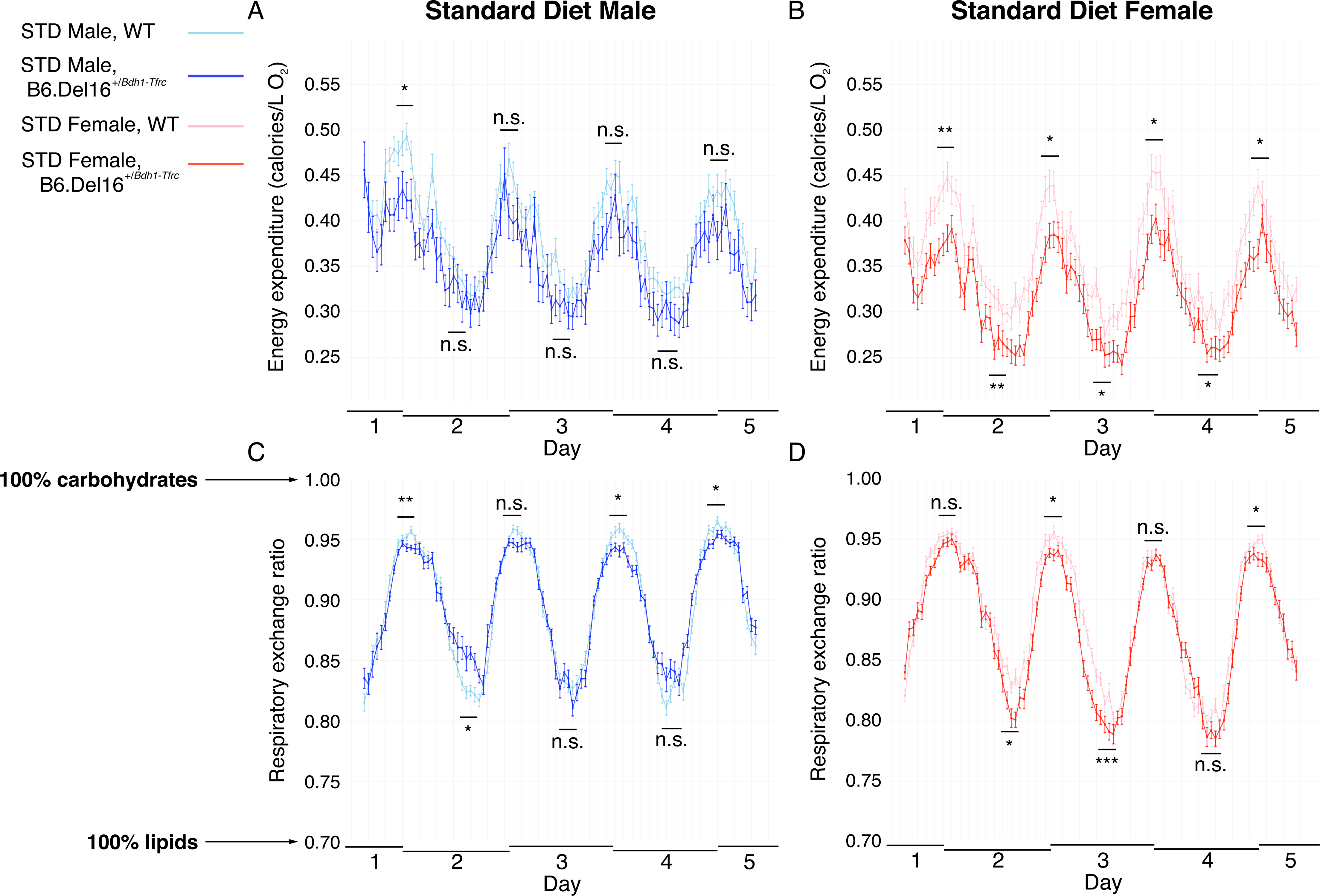
Reduced energy expenditure and respiratory exchange ratio in B6.Del16^+/*Bdh1-Tfrc*^ mice. A and B) Energy expenditure for A) male (n=11 WT, 8 B6.Del16^+/*Bdh1-Tfrc*^) and B) female (n=14 WT, 12 B6.Del16^+/*Bdh1-Tfrc*^) mice on the STD over 5 days in CLAMS/Metabolic Cages. C and D) RER curves for C) male and D) female WT and B6.Del16^+/*Bdh1-Tfrc*^ mice on the STD over 5 days in CLAMS/Metabolic Cages. Data are represented as mean ± SEM. *, p<0.05; **, p<0.01; ***, p<0.001 Statistical analysis was performed using generalized linear models.

To determine whether the weight deficit in B6.Del16^+/*Bdh1-Tfrc*^ mice is simply due to decreased caloric intake, we evaluated food consumption. Food consumption was not significantly different between male or female WT and B6.Del16^+/*Bdh1-Tfrc*^ animals after controlling for the reduced weight of B6.Del16^+/*Bdh1-Tfrc*^ animals (Figure S1A). There were only minor differences in locomotor activity, with male B6.Del16^+/*Bdh1-Tfrc*^ mice *less* active than WT littermates, and no differences between female WT and B6.Del16^+/*Bdh1-Tfrc*^ mice (Figure S1C-D). These data show that the weight deficit in B6.Del16^+/*Bdh1-Tfrc*^ mice is not attributable to a behavioral phenotype such as a decrease in energy consumption or an increase in energy expenditure. Further, the reduced RER in B6.Del16^+/*Bdh1-Tfrc*^ mice indicates that B6.Del16^+/*Bdh1-Tfrc*^ animals are preferentially using lipids as an energy source rather than carbohydrates.

Together, these data support our hypothesis of altered metabolic function associated with the 3q29 deletion and suggest that the 3q29 deletion may be associated with small molecule shifts in the global metabolic environment.

### Untargeted metabolomics reveals small molecule alterations in B6.Del16^+/*Bdh1-Tfrc*^ mice that are highly sex-dependent

To identify metabolic pathway differences between B6.Del16^+/*Bdh1-Tfrc*^ mice and WT littermates, we performed untargeted metabolomic profiling on liver samples using liquid chromatography coupled to ultra-high-resolution mass spectrometry (Go et al., 2015). Males and females were analyzed separately. We compared the lists of all nominally significant metabolic features between WT and B6.Del16^+/*Bdh1-Tfrc*^ samples in the male and female datasets and found that only 22 features were shared (Figure 3A, full details in Supplement), highlighting the sex-dependent effect of the 3q29 deletion on the metabolic environment. Using the top 250 ranked metabolic features, we were able to cluster WT and B6.Del16^+/*Bdh1-Tfrc*^ samples with 84.2%±0.2% accuracy for males (p=0.003) and 76.9%±0% accuracy for females (p=0.006, Figure 3B-E), indicating a strong effect of the 3q29 deletion on the metabolic environment. We also created random forest classifiers using the top 250 ranked features; our male classifier attained an AUC of 0.909±0.002 (p=0.001) and a prediction accuracy of 84.2%±0.2% (p=0.003, Figure 3D), which translates to a 90.9% chance that the classifier will assign an unknown male sample to the correct genotype, and that 84.2% of input samples were correctly classified as WT or B6.Del16^+/*Bdh1-Tfrc*^. Our female classifier attained an AUC of 0.905±0.001 (p=0) and a prediction accuracy of 76.9%±0% (p=0.006, Figure 3E), which translates to a 90.5% chance that the classifier will assign an unknown female sample to the correct genotype, and that 76.9% of input samples were correctly classified as WT or B6.Del16^+/*Bdh1-Tfrc*^. These data show that there is a substantial effect of the 3q29 deletion on the metabolic environment, and this effect is highly sex dependent.

**Figure 3.**
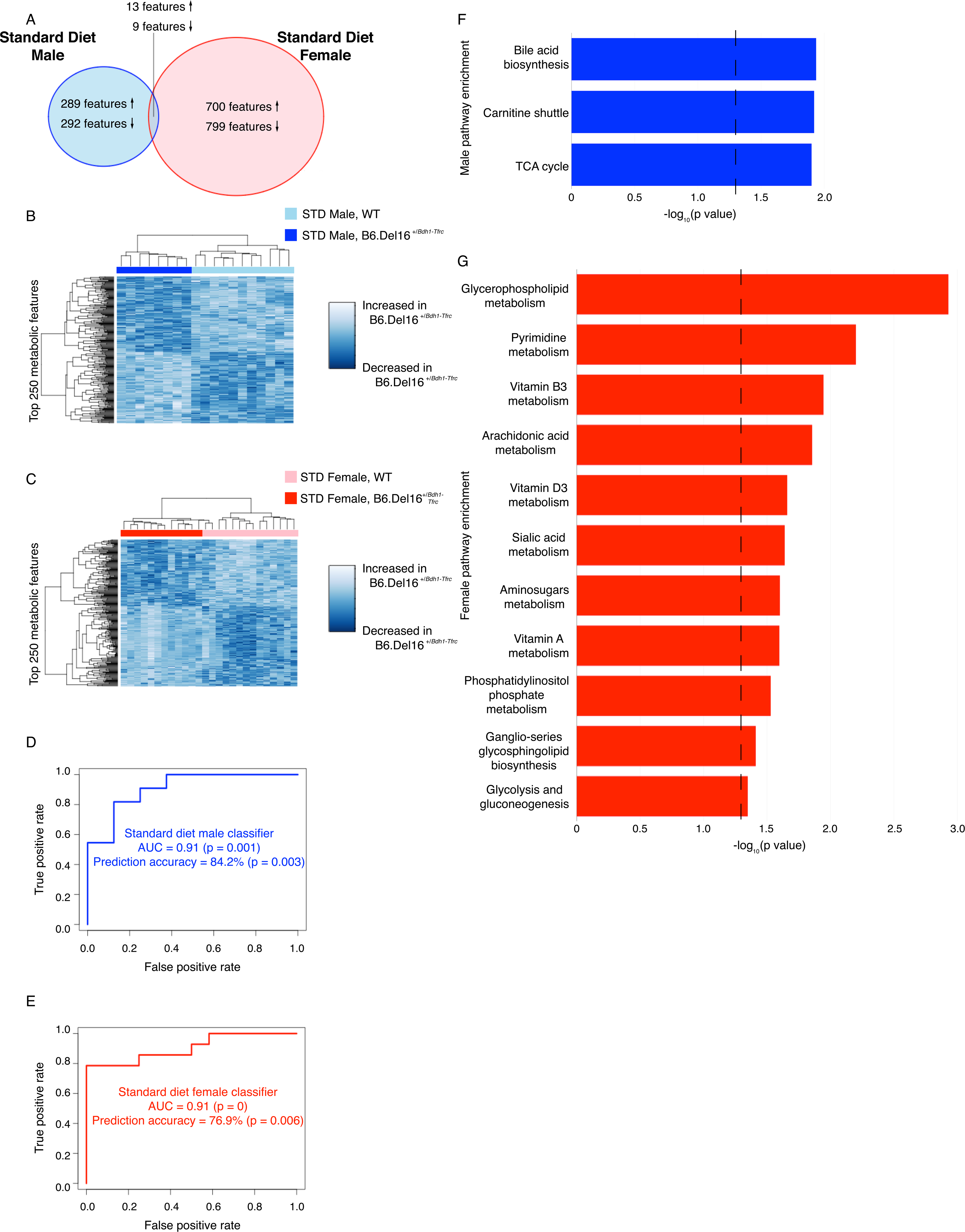
Untargeted metabolomics reveals small molecule alterations in B6.Del16^+/*Bdh1-Tfrc*^ mice that are highly sex-dependent. **A)** Comparison of all nominally significant metabolomic features between the male and female datasets. Up arrows indicate metabolites significantly upregulated in B6.Del16^+/*Bdh1-Tfrc*^ samples, down arrows indicate metabolites significantly downregulated in B6.Del16^+/*Bdh1-Tfrc*^ samples. Also refer to Supplement. B and C) Hierarchical clustering of B) male (n=11 WT, 8 B6.Del16^+/*Bdh1-Tfrc*^) and C) female (n=14 WT, 12 B6.Del16^+/*Bdh1-Tfrc*^) samples using the top 250 ranked metabolomic features. D and E) ROC curves for random forest predictors generated using the top 250 ranked metabolomic features in D) male and E) female datasets. F and G) Altered pathways in B6.Del16^+/*Bdh1-Tfrc*^ mice identified via pathway enrichment analysis of F) male and G) female datasets. Dashed line denotes statistical significance. Statistical significance of ROC curves (D and E) was assessed using 10,000 permutations of the data.

In addition to the lack of individual feature overlap, pathway enrichment analysis of altered features using Mummichog (Li et al., 2013) revealed no pathway-level overlap between the sexes, again highlighting the sex-dependent effects of the 3q29 deletion on metabolism. However, pathways generally related to fat metabolism were identified in both the male and female datasets, including bile acid biosynthesis and the carnitine shuttle in males; and glycerophospholipid metabolism, arachidonic acid metabolism, phosphatidylinositol metabolism, and ganglio-series glycosphingolipid biosynthesis in females (Figure 3F-G). This is consistent with the significant reduction in RER in B6.Del16^+/*Bdh1-Tfrc*^ mice, which indicated a preference for dietary lipids as fuel in B6.Del16^+/*Bdh1-Tfrc*^ mice as compared to WT littermates. These metabolic pathway data further demonstrate that metabolism of dietary fat may be altered in both male and female B6.Del16^+/*Bdh1-Tfrc*^ animals.

### A high-fat diet attenuates the B6.Del16^+/*Bdh1-Tfrc*^ weight deficit and affects RER in a sex-specific manner

From the RER data, we determined that B6.Del16^+/*Bdh1-Tfrc*^ animals preferentially use fat as an energy source; untargeted metabolomics data confirmed there are differences in fat metabolism pathways in B6.Del16^+/*Bdh1-Tfrc*^ mice on a standard diet (STD). As the STD contains only 13.4% of total calories from fat; we hypothesized that increased availability of fat calories would cause B6.Del16^+/*Bdh1-Tfrc*^ animals to gain weight and remedy the 3q29 deletion-associated weight deficit. To test this hypothesis, we implemented a high-fat diet (HFD) challenge from postnatal day 21 to euthanasia (16-20 weeks), using a commercially available diet (TD.88137, Teklad Custom Diets, Envigo) containing 42% of total calories from fat. The weight deficit in B6.Del16^+/*Bdh1-Tfrc*^ mice was partially ameliorated in female mice but was largely unchanged in male mice after HFD treatment (Figure 4A-B). After HFD treatment, male B6.Del16^+/*Bdh1-Tfrc*^ mice on the HFD weighed on average 1.72 g less than WT littermates (p=0.015, Figure 4A), compared to 1.61 g less than WT littermates on the STD (Rutkowski et al., 2019). Female B6.Del16^+/*Bdh1-Tfrc*^ mice on the HFD weighed on average 1.66 g less than WT littermates (p=0.004, Figure 4B), versus 2.24 g less than WT littermates on the STD (Rutkowski et al., 2019). When these effect sizes were considered relative to total body weight, the effect of the HFD intervention became clearer. Male B6.Del16^+/*Bdh1-Tfrc*^ mice on the HFD were 4.33% smaller than WT littermates at 16 weeks, compared to 5.16% smaller on the STD. Female B6.Del16^+/*Bdh1-Tfrc*^ mice on the HFD were 5.16% smaller than WT littermates at 16 weeks, compared to 10.29% smaller on the STD at the same timepoint. These data demonstrate a sex-specific effect of the 3q29 deletion on the response to the HFD, further supporting the differential impact of the 3q29 deletion on metabolism in B6.Del16^+/*Bdh1-Tfrc*^ mice.

**Figure 4.**
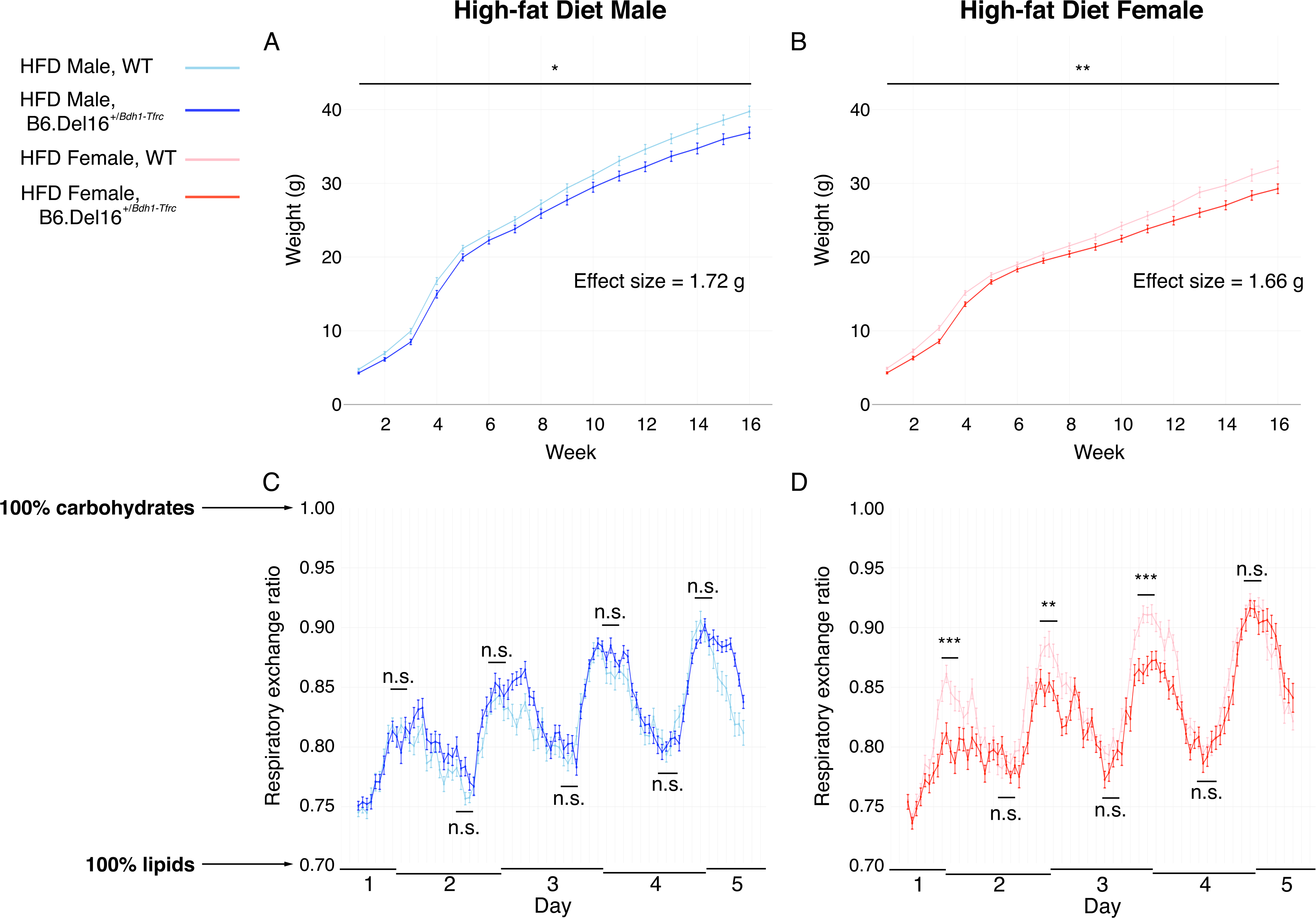
A high-fat diet reduces the B6.Del16^+/*Bdh1-Tfrc*^ weight deficit and affects RER in a sex-specific manner. A and B) 16-week growth curves for HFD-treated A) male (n=50 WT, 30 B6.Del16^+/*Bdh1-Tfrc*^) and B) female (n=42 WT, 32 B6.Del16^+/*Bdh1-Tfrc*^) mice. C) Raw effect size of the 3q29 deletion on weight in HFD-treated male and female mice. D) Effect size of the 3q29 deletion on weight relative to average WT body weight at week 16 in HFD-treated male and female mice. E and F) RER curves for E) male (n=10 WT, 10 B6.Del16^+/*Bdh1-Tfrc*^) and F) female (n=10 WT, 10 B6.Del16^+/*Bdh1-Tfrc*^) mice on the HFD over 5 days in CLAMS/Metabolic Cages. Data are represented as mean ± SEM. n.s., p>0.05; *, p<0.05; **, p<0.01; ***, p<0.001 Statistical analysis of growth curves (A-D) was performed using generalized estimating equations. Statistical analysis of RER (E-F) was performed using generalized linear models.

To further investigate the metabolic consequences of the HFD intervention, we performed 5 days of indirect calorimetry on the HFD-treated male and female B6.Del16^+/*Bdh1-Tfrc*^ and WT mice. There were no differences in RER between male WT and B6.Del16^+/*Bdh1-Tfrc*^ mice, indicating that male WT and B6.Del16^+/*Bdh1-Tfrc*^ animals have similar energy source utilization on the HFD (Figure 4C). By contrast, RER peaks were reduced in female B6.Del16^+/*Bdh1-Tfrc*^ mice on the HFD compared to WT animals, consistent with the STD data (Figure 4D). Even after HFD treatment, metabolism in female B6.Del16^+/*Bdh1-Tfrc*^ mice was still shifted toward preferentially utilizing lipids rather than carbohydrates, whereas male B6.Del16^+/*Bdh1-Tfrc*^ mice had comparable macronutrient utilization to WT animals after the HFD intervention. These data suggest that the HFD rescued the shift in macronutrient utilization in males, but females may have a pronounced need for dietary lipids that was not fully satisfied by the HFD intervention. Consistent with the STD results, there were no differences between male or female WT and B6.Del16^+/*Bdh1-Tfrc*^ mice in food or water consumption (Figure S1E-F) or activity (Figure S1G-H). There was no difference in energy expenditure in male HFD animals (Figure S1I) and a slight difference in energy expenditure in female HFD animals (Figure S1J). Together, these data provide further support to the conclusion that male and female B6.Del16^+/*Bdh1-Tfrc*^ mice are differentially impacted by 3q29 deletion-associated metabolic deficits.

### Widespread changes in the global metabolic environment of B6.Del16^+/*Bdh1-Tfrc*^ mice after HFD treatment

We found that the HFD intervention altered RER in male B6.Del16^+/*Bdh1-Tfrc*^ mice and partially ameliorated the 3q29 deletion-associated weight deficit in female B6.Del16^+/*Bdh1-Tfrc*^ mice. Based on these data, we hypothesized that the HFD intervention may also affect changes in the global metabolic environment of B6.Del16^+/*Bdh1-Tfrc*^ animals. To test this hypothesis, we performed untargeted metabolomic profiling of liver samples from HFD-treated animals (Go et al., 2015). Males and females were analyzed separately. Similar to the results from our metabolomics analysis of STD-treated animals, comparison of all nominally significant metabolic features between the male and female datasets revealed only 7 shared features between sexes (Figure 5A, full details in Supplement), highlighting a substantial sex-dependent effect of the 3q29 deletion on the metabolic environment after HFD treatment. Using the top 250 ranked metabolites, WT and B6.Del16^+/*Bdh1-Tfrc*^ samples clustered with 100%±0% accuracy in males (p=0) and 95%±0% accuracy in females (p=0, Figure 5B-E). These data suggest that even after the HFD intervention, pronounced metabolic differences exist between the WT and B6.Del16^+/*Bdh1-Tfrc*^ mice. Additionally, random forest classifiers using the top 250 ranked features achieved excellent classification (AUC=1±0 in males and 1±0.001 in females) and high prediction accuracy (100%±0% in males and 95%±0% in females, Figure 5D-E) in both sexes. Pathway enrichment analysis of altered features using Mummichog (Li et al., 2013) identified pathways with diverse functions in both datasets, including pyrimidine metabolism and aspartate and asparagine metabolism in males, and vitamin B6 metabolism and propanoate metabolism in females (Figure 5F-G). Pathways related to fat metabolism were identified in the female dataset, including *de novo* fatty acid biosynthesis and ganglio-series glycosphingolipid biosynthesis, as they were in the STD pathway analysis. This result is concordant with our finding of persistent RER shifts in female, but not male, B6.Del16^+/*Bdh1-Tfrc*^ mice. At the small molecule level, the HFD treatment did not restore fat metabolism functions in female B6.Del16^+/*Bdh1-Tfrc*^ mice to WT levels, supporting our hypothesis that the HFD intervention did not fully satisfy the increased metabolic demand for fat in female B6.Del16^+/*Bdh1-Tfrc*^ mice. As seen in the STD result and the lack of individual feature overlap in the HFD datasets, there was no pathway-level overlap between the sexes, further demonstrating that a robust sex effect of the 3q29 deletion on the metabolic environment remains after HFD treatment.

**Figure 5.**
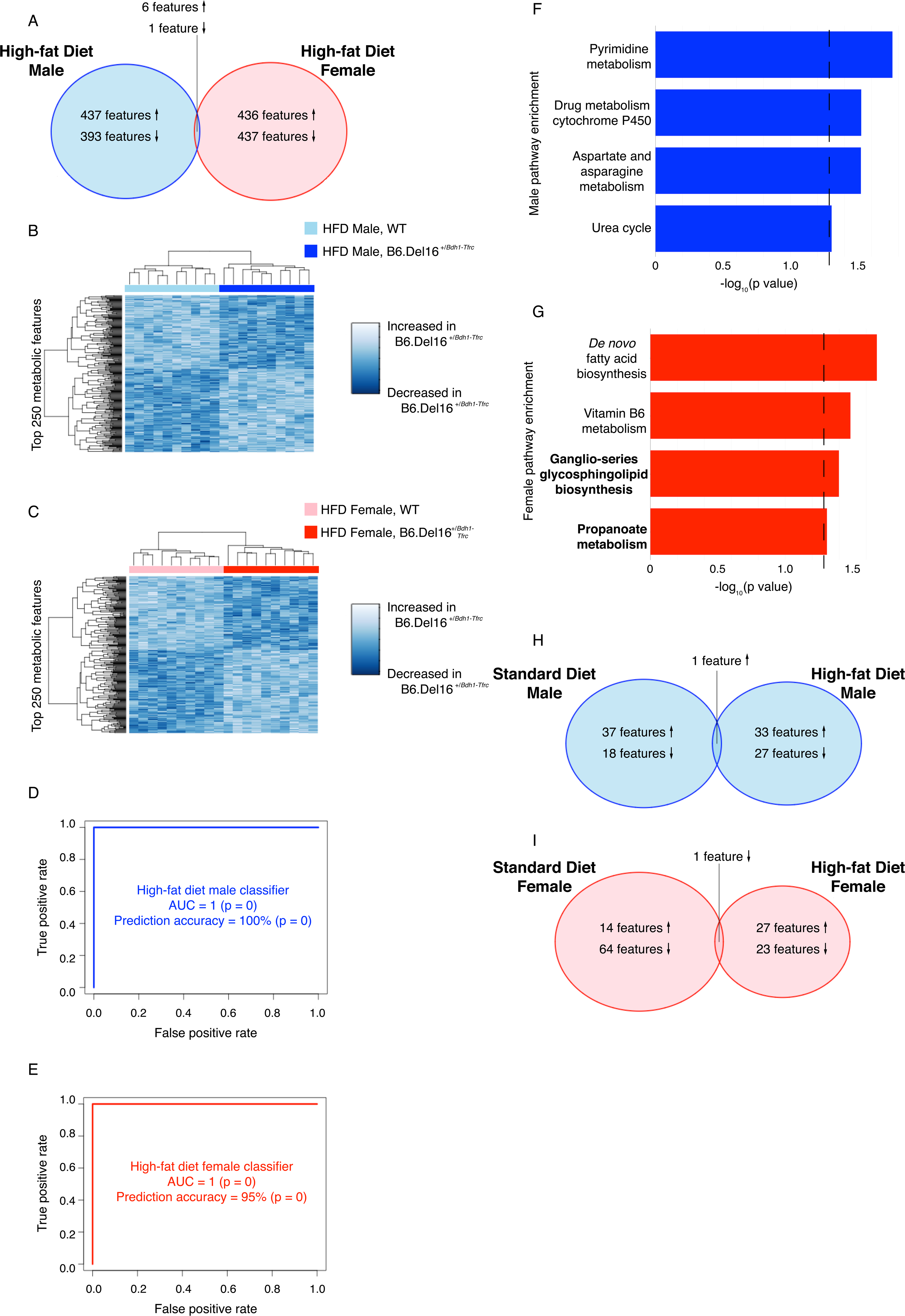
Widespread changes in the global metabolic environment of B6.Del16^+/*Bdh1-Tfrc*^ mice after HFD treatment. A) Comparison of all nominally significant metabolomic features between the HFD-treated male and female datasets. Up arrows indicate metabolites significantly upregulated in B6.Del16^+/*Bdh1-Tfrc*^ samples, down arrows indicate metabolites significantly downregulated in B6.Del16^+/*Bdh1-Tfrc*^ samples. Also refer to Supplement. B and C) Hierarchical clustering of HFD-treated B) male (n=10 WT, 10 B6.Del16^+/*Bdh1-Tfrc*^) and C) female (n=10 WT, 10 B6.Del16^+/*Bdh1-Tfrc*^) samples using the top 250 ranked metabolomic features. D and E) ROC curves for random forest predictors generated using the top 250 ranked metabolomic features in HFD-treated D) male and E) female datasets. F and G) Altered pathways in HFD-treated B6.Del16^+/*Bdh1-Tfrc*^ mice identified via pathway enrichment analysis of F) male and G) female datasets. Dashed line denotes statistical significance. Bold text denotes pathways that were identified in both the STD and HFD experiments. H and I) Comparison of nominally significant annotated features between STD-treated and HFD-treated H) male and I) female datasets. Up arrows indicate metabolites significantly upregulated in B6.Del16^+/*Bdh1-Tfrc*^ samples, down arrows indicate metabolites significantly downregulated in B6.Del16^+/*Bdh1-Tfrc*^ samples. Also refer to Supplement. Statistical significance of ROC curves (D and E) was assessed using 10,000 permutations of the data.

To test for changes in metabolism resulting from the HFD intervention, we performed a direct comparison between the metabolomics results from the STD and HFD cohorts, stratified by sex. We compared the statistically significant high-confidence annotated metabolic features in each dataset. In the male datasets, there were 61 significant annotated features in the STD cohort, and 56 significant annotated features in the HFD cohort, and only four of those features were identified in both datasets (Figure 5H, full details in Supplement). The comparison between the female STD and HFD cohorts yielded similar results; there were 79 significant annotated features in the female STD cohort and 51 significant annotated features in the female HFD cohort, with only three features identified in both datasets (Figure 5I, full details in Supplement). Further, we found that these major shifts in the metabolic environment were recapitulated at the pathway level; ganglio-series glycosphingolipid biosynthesis was the only metabolic pathway identified in both female cohorts, and pyrimidine metabolism was identified in the female STD cohort and the male HFD cohort (Figure 5G). Together, these data demonstrate that that the HFD intervention resulted in major shifts in the global metabolic environment of both male and female animals, but response to the HFD was highly sex-specific. Further, the finding that ganglio-series glycosphingolipid biosynthesis was identified in both female cohorts suggests that the HFD intervention did not fully restore fat metabolism in female B6.Del16^+/*Bdh1-Tfrc*^ mice to WT levels. Finally, although body weight of B6.Del16^+/*Bdh1-Tfrc*^ mice on the HFD approached that of WT littermates, the substantial genotype differences in the global metabolic environment show that the underlying metabolism of B6.Del16^+/*Bdh1-Tfrc*^ mice still did not recapitulate that of WT animals.

### HFD treatment does not affect B6.Del16^+/*Bdh1-Tfrc*^ brain size or behavioral phenotypes

Reduced brain size has been described in both mouse models of the 3q29 deletion (Baba et al., 2019; Rutkowski et al., 2019). Additionally, prior work by our team identified an increased brain:body weight ratio in female, but not male, B6.Del16^+/*Bdh1-Tfrc*^ mice compared to WT littermates (Rutkowski et al., 2019). An increase in the brain:body weight ratio has been observed in human and animal models of starvation, lending support to the hypothesis that the brain is metabolically privileged (Addis et al., 1936; Jackson, 1915; Keys et al., 1950; Schmidt et al., 1945). Because female B6.Del16^+/*Bdh1-Tfrc*^ mice showed metabolic improvement after HFD treatment, we hypothesized that the brain:body weight ratio in female B6.Del16^+/*Bdh1-Tfrc*^ mice on the HFD would be reduced to WT levels. We found that, consistent with prior reports, brain weight was reduced in both male (p=3E-6) and female (p=0.04) B6.Del16^+/*Bdh1-Tfrc*^ mice relative to WT littermates (Figure 6A). Additionally, we found that the brain:body weight ratio in male B6.Del16^+/*Bdh1-Tfrc*^ mice was identical to that in WT animals (p=1); however, the brain:body weight ratio in female B6.Del16^+/*Bdh1-Tfrc*^ mice was increased relative to WT littermates (p=0.04, Figure 6B). These data show that while the HFD intervention resulted in metabolic changes in male and female B6.Del16^+/*Bdh1-Tfrc*^ mice, and partially ameliorated the weight deficit in female B6.Del16^+/*Bdh1-Tfrc*^ mice, these positive effects did not extend to the brain:body weight ratio, indicating that early neurodevelopmental processes may not have been impacted by the HFD.

**Figure 6.**
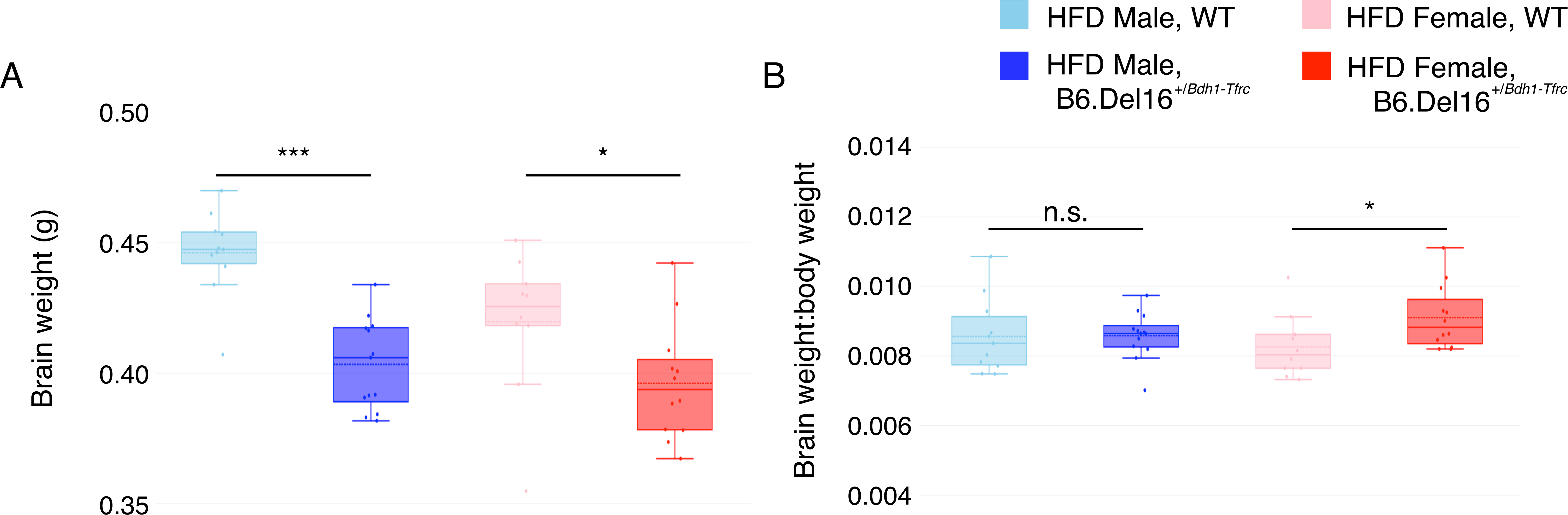
HFD treatment does not affect B6.Del16^+/*Bdh1-Tfrc*^ brain size. **A)** Brain weight in HFD-treated male (n=10 WT, 10 B6.Del16^+/*Bdh1-Tfrc*^) and female (n=10 WT, 10 B6.Del16^+/*Bdh1-Tfrc*^) mice. **B)** Brain weight:body weight ratio in HFD-treated male and female mice. For each box plot, the dashed line indicates the mean value, and the solid line indicates the median. n.s., p>0.05; *, p<0.05; ***, p<0.001 Statistical analysis was performed using unpaired t-tests.

The 3q29 deletion has a well-established association with neurodevelopmental and neuropsychiatric phenotypes (Ballif et al., 2008; Cox and Butler, 2015; Itsara et al., 2009; Kirov et al., 2012; Marshall et al., 2017; Mulle, 2015; Mulle et al., 2010; Sanders et al., 2015; Szatkiewicz et al., 2014; Willatt et al., 2005); behavioral deficits have also been identified in two independent mouse models of the 3q29 deletion (Baba et al., 2019; Rutkowski et al., 2019). To understand the effect of the HFD intervention on behavioral phenotypes in B6.Del16^+/*Bdh1-Tfrc*^ mice, we performed a pilot study using a battery of assays designed to test learning and memory, acoustic startle, sensorimotor gating, and amphetamine sensitivity. We replicated several phenotypes, including spatial learning and memory deficits, an elevated acoustic startle response, sensorimotor gating deficits, and attenuated amphetamine-induced locomotion (Rutkowski et al., 2019) (full details in Supplement). We observed some sex differences between our findings and previously published results (Rutkowski et al., 2019); however, the present study focused on diet and was sufficiently powered for metabolic analyses but not for subtle behavioral phenotypes.

There was a significant main effect of the diet intervention (p<0.05) in the Morris water maze (MWM), acoustic startle, fear conditioning, and amphetamine-induced locomotor activity for both male and female B6.Del16^+/*Bdh1-Tfrc*^ animals (full details in Supplement). However, the effect of the diet was shared across genotypes; the HFD intervention did not differentially impact behavior based on genotype (full details in Supplement). Together, these data demonstrate that the HFD did not introduce any appreciable changes to behavioral phenotypes of B6.Del16^+/*Bdh1-Tfrc*^ mice, suggesting that 3q29 deletion-related metabolic and behavioral phenotypes may arise from uncoupled, independent mechanisms.

## DISCUSSION

This is the first study to examine metabolic function associated with the 3q29 deletion in a comprehensive manner. Previous work by our group characterized a persistent, sex-dependent weight deficit in B6.Del16^+/*Bdh1-Tfrc*^ mice (Rutkowski et al., 2019), and a study of an independently generated mouse model of the 3q29 deletion also identified a reduced weight phenotype in males, but did not examine female animals (Baba et al., 2019). In the current study we expanded upon this work by exploring mechanisms that may lead to this reduced weight phenotype, incorporating both male and female animals in our study design to specifically assess the effect of sex on metabolic phenotypes (Figure 1). We identified pervasive sex effects of the 3q29 deletion on metabolic function, including differential effects on the metabolic environment and the response to HFD. After HFD treatment, female animals showed a change in weight, and male and female animals showed contrasting changes in the metabolic profile. The HFD intervention did not rescue behavioral phenotypes in male or female animals, suggesting that metabolic and behavioral phenotypes in the context of the 3q29 deletion may arise from independent mechanisms. This study is a step toward unraveling the biology underlying the development of diverse phenotypic outcomes in 3q29 deletion syndrome. Furthermore, the substantial sex effects identified here show it is imperative to evaluate sex as a biological variable in metabolic studies.

In light of our data, in particular the RER shifts in STD-treated animals and the response to the HFD, we conclude that B6.Del16^+/*Bdh1-Tfrc*^ mice preferentially use lipids as an energy source. This preference for lipids was more pronounced in female animals and was partially corrected by the HFD. The HFD intervention also affected substantial changes in the global metabolic environment as assessed via untargeted metabolomics; the small molecule profile was substantially altered after HFD treatment in both males and females. Consistent with the lack of overlap in the small molecule profile after the HFD intervention, the altered metabolic pathways in B6.Del16^+/*Bdh1-Tfrc*^ mice had minimal overlap between the STD and HFD datasets. Notably, the ganglio-series glycosphingolipid biosynthesis pathway was altered in both STD- and HFD-treated female B6.Del16^+/*Bdh1-Tfrc*^ mice; this supports our conclusion that the HFD did not fully rescue fat metabolism deficits in B6.Del16^+/*Bdh1-Tfrc*^ females and further emphasizes the persistent sex-dependent effects of the 3q29 deletion.

The striking lack of overlap in male and female metabolic effects of the 3q29 deletion was unexpected. We sought to integrate our data with the larger metabolomics literature, but our initial literature search suggested that most studies did not include sex-stratified analysis of mouse metabolomics data. Our literature search also indicated that female mice were often excluded from the study design. To test this hypothesis, we conducted a formal meta-analysis of the literature. We searched for the terms “mouse metabolomics” and identified 2,601 possible studies (full details in Supplement). We selected 500 of these studies at random and classified them according to whether both sexes were included, and whether sex-stratified analysis was conducted. Of the 500 publications we evaluated, only 44 (8.8%) included both sexes, and only 17 (3.4%) analyzed the data separately by sex. Of these 17 studies, 9 (52.9%) reported substantial sex-dependent differences in the metabolome, indicating there may be widespread but unappreciated differences between male and female mouse metabolomics data. Our conclusion that the metabolomic differences due to the 3q29 deletion are highly-sex dependent is consistent with the larger, albeit limited, scope of the literature. This literature meta-analysis highlights a pronounced knowledge gap in the field of metabolomics research, as there may be substantial, unappreciated sex-dependent metabolic differences in mouse models. These results have profound implications for the design of future metabolic studies; it is imperative that males and females be included and analyzed separately to rigorously assess the role of sex as a biological variable.

There are well-established links between sex and metabolism. Males and females have different patterns of fat deposition, and differences in fat metabolism have been identified in both humans and rodents (Blaak, 2001; Faulds et al., 2012; Mayes and Watson, 2004; Ockner et al., 1979; Soler-Argilaga and Heimberg, 1976; Wu and O’Sullivan, 2011). Studies in rodents have revealed that the sex chromosome complement affects fat metabolism; methods such as the four core genotypes model (De Vries et al., 2002) have helped to disentangle the effects of sex hormones and sex chromosomes on fat metabolism (Chen et al., 2012; Chen et al., 2013; Link et al., 2013; Link and Reue, 2017). These findings are supported at the level of gene expression. A large proportion of liver-expressed genes in humans show sex-biased expression, and the complement of sex-biased genes are enriched for fat metabolism functions (Zhang et al., 2011). Sex hormones, specifically estrogen, also appear to have a role in sex-dependent differences in fat metabolism; oral estrogen therapy in postmenopausal women leads to well-documented changes in fat metabolism (O’Sullivan et al., 1998; Walsh et al., 1991), and endogenous levels of sex hormones also impact fat metabolism and fat distribution (Faulds et al., 2012; Link and Reue, 2017; Mayes and Watson, 2004; Ockner et al., 1980; Palmisano et al., 2018; Wu and O’Sullivan, 2011). Data from animal models have revealed pervasive roles for estrogen and estrogen-related signaling in metabolic processes including fat metabolism and storage (Faulds et al., 2012). An independent study by our group using transcriptome network analysis identified a co-expression module significantly enriched for estrogen-dependent signaling (p=5.08E-06) that contained the 3q29 deletion interval gene *PAK2* (Sefik et al., 2020). Together with the existing literature on sex differences in fat metabolism and our finding that male and female B6.Del16^+/*Bdh1-Tfrc*^ mice are differentially affected by 3q29 deletion-associated metabolic phenotypes, this finding suggests that sex is an important consideration in defining the biological mechanisms underscoring phenotypes in 3q29del.

The links between the 3q29 deletion and metabolic function (Baba et al., 2019; Glassford et al., 2016; Rutkowski et al., 2019) are not unique in the broader context of recurrent CNV disorders. Weight changes and failure to thrive are associated with many recurrent CNVs, including the 22q11.2, 16p11.2, 17p11.2, and 1q21.1 loci (Arbogast et al., 2016; Bochukova et al., 2010; Burns et al., 2010; Edelman et al., 2007; Giardino et al., 2014; Jacquemont et al., 2011; Lacaria et al., 2012; Matthiesen et al., 2016; Mefford et al., 2008; Pettersson et al., 2017; Smith et al., 2002; Voll et al., 2017; Walters et al., 2010; Zufferey et al., 2012). Evidence has shown that pediatric feeding disorders and nutrient deficiencies can exacerbate existing neurodevelopmental and cognitive deficits (O’Brien et al., 1991; Silverman and Tarbell, 2009; Volkert et al., 2016). In this context, addressing feeding disorders and metabolic concerns in individuals with CNV disorders should be a priority, to minimize the adverse effects of poor nutrition on long-term outcomes. In the present study, we found that a HFD treatment improved metabolic phenotypes but did not affect behavioral phenotypes in B6.Del16^+/*Bdh1-Tfrc*^ mice, suggesting that a lipid-rich dietary intervention in humans with 3q29del may improve weight phenotypes and nutritional status without exacerbating behavioral phenotypes.

The effects of recurrent CNVs on growth-related phenotypes have been relatively well-described; however, the current understanding of the biological mechanisms leading to these phenotypes is lacking. Recent molecular studies have started to elucidate these mechanisms for 22q11.2 deletion syndrome; mitochondrial dysfunction has been identified as a key contributor to neuronal and synaptic defects associated with the deletion (Fernandez et al., 2019; Gokhale et al., 2019; Napoli et al., 2015; Wesseling et al., 2017). Additionally, a recent study of a mouse model of the 16p11.2 deletion and duplication revealed opposite effects of the CNV on metabolic function (Arbogast et al., 2016). However, these studies largely focused on targeted metabolic measurements, and failed to address sex as a potential mediator of metabolic phenotypes. The incorporation of untargeted approaches into studies of CNV disorders, and the rigorous interrogation of the role of sex in CNV-associated phenotypes, may expedite our understanding of the biological mechanisms at play in these complex disorders.

While there are established links between neurodevelopmental and neuropsychiatric disorders and metabolic function, we found that the HFD intervention improved metabolic phenotypes but did not affect behavioral phenotypes in B6.Del16^+/*Bdh1-Tfrc*^ mice. There are several possible explanations for this outcome. First, the HFD intervention was implemented starting at postnatal day 21. It is possible this was too late to impact the neurodevelopmental processes that contribute to 3q29 deletion phenotypes. In future experiments, the HFD could be applied to pregnant or nursing dams, potentially exposing 3q29 deletion pups to abundant lipid sources earlier in development (Purcell et al., 2011). It is also possible that the behavioral assays we used and/or the sample size we evaluated could only detect large effects on behavior; the HFD may have caused subtle behavioral improvements that we were unable to detect with the behavioral battery we performed. Additionally, the HFD intervention only targeted fat metabolism, while other metabolic alterations in B6.Del16^+/*Bdh1-Tfrc*^ mice would not have been improved by the HFD. Our observation that the weight deficit was only partially ameliorated in B6.Del16^+/*Bdh1-Tfrc*^ mice supports this hypothesis, and suggests that the underlying biology of the 3q29 deletion involves multiple metabolic processes. Data from the present study suggest that metabolism and neurodevelopment may be unlinked in the context of the 3q29 deletion and may be influenced by separate sets of genes within the deletion interval.

The present study was the first to identify metabolic deficits in the context of the 3q29 deletion. Furthermore, we found pervasive sex-specific effects of the 3q29 deletion; these findings are supported by transcriptome network analysis by our team that identified a module enriched for estrogen-related signaling (Sefik et al., 2020). These results have important implications, both for the 3q29 deletion specifically and for metabolic and mechanistic studies more generally. Our findings suggest that metabolic and behavioral phenotypes may arise from independent mechanisms in the context of the 3q29 deletion, and that these mechanisms may be sex specific. This study underscores a critical need for metabolic and mechanistic experiments to include samples from both male and female subjects, and to analyze the data in a sex-specific manner. Due to the substantial, well-documented metabolic, medical, and neurodevelopmental and neuropsychiatric differences between males and females (Abel et al., 2010; Kamitaki et al., 2020; Khramtsova et al., 2019; Mandy et al., 2012; Ngo et al., 2014; Regitz-Zagrosek and Kararigas, 2017; Wu and O’Sullivan, 2011), it is not surprising that by analyzing only one sex, or by pooling data from males and females, important metabolic insights may be obscured.

Additionally, mechanistic studies in complex disorders that combine data from males and females may miss important sex dependent differences in mechanism, which could delay advancements in available therapeutics. Together, our data highlight sex dependent differences in metabolic function in a mouse model of the 3q29 deletion, adding to our current understanding of 3q29del and creating a framework for future mechanistic studies of complex disorders.

## Supporting information

Key Resources Table

## ACKNOWLEDGEMENTS

The work was supported by NIH 1R01MH110701-01A1, NIH R01GM097331, NIH R56MH116994, NIH T32 GM0008490, and the Emory University Treasure Your Exceptions Project. This study was supported in part by the Mouse Transgenic and Gene Targeting Core and the Rodent Behavioral Core, which are subsidized by the Emory University School of Medicine and are part of the Emory Integrated Core Facilities. The authors acknowledge the contributions of the members of the Emory 3q29 Project: Hallie Averbach, Emily Black, T. Lindsey Burrell, Grace Carlock, Joseph F. Cubells, David Cutler, Roberto Espana, Michael J. Gambello, Katrina Goines, Henry R. Johnston, Cheryl Klaiman, Sookyong Koh, Elizabeth J. Leslie, Longchuan Li, Bryan Mak, Trenell Mosley, Melissa Murphy, Derek Novacek, Rossana Sanchez, Celine A. Saulnier, Jason Schroeder, Esra Sefik, Brittney Sholar, Sarah Shultz, Nikisha Sisodiya, Steven Sloan, Elaine F. Walker, and Zhexing Wen.

## AUTHOR CONTRIBUTIONS

RMP performed the research, performed the statistical analysis, produced all figures and tables, and wrote the manuscript. RHP, TPR, TM, KP, and MRS helped perform the research. MPE helped with statistical analyses and interpretation. PAD, MRS, DPJ, STW, MEZ, TC, DW, and JGM provided guidance on analyzing and interpreting data. GJB, STW, MEZ, TC, DW, and JGM edited the manuscript. GJB, STW, TC, DW, and JGM were the principal investigators responsible for study direction. All authors participated in commenting on the drafts and have read and approved the final manuscript.

## STAR METHODS

### Resource Availability

#### Lead Contact

Further information and requests for resources and reagents should be directed to and will be fulfilled by the Lead Contact, Jennifer Mulle.

#### Materials Availability

This study did not generate new unique reagents.

#### Data and Code Availability

The code supporting the current study is available from the corresponding author upon reasonable request. The datasets supporting the current study are publicly available at http://dx.doi.org/10.17632/g3v2nt657z.1.

### Experimental Model and Subject Details

#### Animals, Husbandry, and Diets

All studies were performed on male and female C57BL/6N-Del16^+/*Bdh1-Tfrc*^ (B6.Del16^+/*Bdh1-Tfrc*^, MGI:6241487) mice and wild type (WT) littermates (Rutkowski et al., 2019). All animals were maintained on a C57BL/6N background sourced from Charles River Laboratories. Mice were group housed (maximum of 5 animals per cage) during the entire experiment, except for a five-day separation during indirect calorimetry when a subset of mice were singly housed. Mice were on a 12-hour light/dark cycle and were given food and water *ad libitum*. Starting at postnatal day 21, mice were fed either a standard diet (STD, LabDiet 5001) low in fat (13.4% energy from fat) or a high-fat diet (HFD, Teklad TD.88137, 42.0% energy from fat) for the remainder of their lives. Body weight was monitored weekly from 1-16 weeks of age. Indirect calorimetry and behavioral assays were performed on mice between 16-20 weeks of age. At the conclusion of indirect calorimetry, mice were euthanized, and tissues were collected for metabolomics analysis. Mice were not fasted prior to euthanasia and tissue collection. All animal protocols were performed under the approved guidelines of the Emory University Institutional Animal Care and Use Committee. Both males and females were used in all experiments, and the data were analyzed separately. Number of animals used in experiments is indicated in figure legends.

### Method Details

#### Indirect Calorimetry

Mouse metabolic rate was assessed by indirect calorimetry for 5 days in Oxymax chambers using the Comprehensive Lab Animal Monitoring System (Oxymax CLAMS-HC, Columbus Instruments). Mice were singly housed with *ad libitum* access to water and food and were maintained at 20-22°C under a 12/12 hr light/dark cycle (light period 07:00-19:00). A mass-sensitive flow meter was used to maintain a constant airflow of 0.6 L/min. n=8-12 mice/group; age=16-18 weeks

#### Metabolomics

Untargeted metabolomics analysis on mouse liver tissue was performed as previously described (Go et al., 2015). Briefly, supernatants were analyzed by liquid chromatography coupled to ultra-high-resolution mass spectrometry (LC-HRMS). A quality control pooled reference sample (QStd3) was included at the beginning and end of each analytical batch of 20 samples for quality control and quality assurance (Go et al., 2015). Samples were analyzed in triplicate by liquid chromatography with Fourier transform mass spectrometry (Dionex, Ultimate 3000, Velos, Thermoe Fisher) with hydrophilic interaction liquid chromatography (HILIC) with positive electrospray ionization (ESI+) mode and reverse phase (C18) chromatography with positive electrospray ionization (ESI+) mode and resolution of 70,000 (Jones et al., 2016). Four blind replicate samples were included in the STD analysis (Figure S2A-H) and two blind replicate samples were included in the HFD analysis (Figure S2I-L) to ensure data quality. Spectral *m/z* features were acquired in scan range 85-1,250 mass-to-charge ratio (*m/z*). Raw data files were extracted using apLCMSv6.3.3 (Yu et al., 2013) with xMSanalyzer v2.0.7 (Uppal et al., 2013), followed by batch correction with ComBat (Johnson et al., 2007). Resulting mass spectrometry data, referred to as *m/z* features, included accurate mass *m/z*, retention time (s), and ion abundance. All samples were analyzed in triplicate, and the feature intensities were median summarized. n=8-12 mice/group; age=17-19 weeks

#### Behavior Tests

##### Morris Water Maze

The Morris water maze (MWM) was conducted to test for deficits in spatial learning and memory. Briefly, the MWM was conducted in a circular tank (52 inches in diameter) filled with water, made opaque with white paint, at 23°C. A hidden circular platform (30 cm in diameter) in the northwestern quadrant of the tank was present 1 cm below the surface of the water. The tank was surrounded by white walls on the north and east sides and white curtains on the west and south sides, all containing external cues for spatial reference. Mice were trained to find the hidden platform over 5 days by being released into the tank from each quadrant (north, south, east, and west) in a randomized order each day. Each trial lasted a maximum of 60 s; if a mouse did not find the platform in that time, it was guided to the platform and allowed to rest on the platform for 10 s. On the sixth day, the platform was removed from the tank and a probe trial was conducted, in which the mouse was placed in the tank at the south start point and allowed to swim for 60 s. An automated tracking system (TopScan, CleverSys) was used during training to record the latency and distance to the platform and swim speed and was used during the probe trial to record the duration and distance the mice spent in each quadrant of the maze. n=5-13 mice/group; age=16-20 weeks

##### Acoustic Startle and Prepulse Inhibition

To test for deficits in the acoustic startle response and sensorimotor gating, acoustic startle and prepulse inhibition (PPI) were performed. To test the acoustic startle response, mice were subjected to a series of increasing startle tons (75, 80, 85, 90, 95, 100, 115, and 120 db) for 40 ms each, and the response of the animal was measured by the SR-LAB startle response system (San Diego Instruments) accelerometer. A startle curve was constructed to ensure the animal was responding to the increasing stimulus. On the second day, PPI was evaluated. The mice were exposed to 6 blocks of startle conditions with each block consisting of 12 trials, so that each trial was presented to the animal 6 times. The 12 trials were randomly ordered in each block, and the animal’s response to the stimulus was measured after each trial. The 12 trials consisted of the following conditions: background (68 db) for 20 ms; startle (120 db) for 40 ms; prepulses 1-5 (PP1=70 db, PP2=74 db, PP3=78 db, PP4=82 db, PP5=86 db) for 20 ms each; and the prepulse.startle combinations (PP1.startle, PP2.startle, PP3.startle, PP4.startle, and PP5.startle), where each prepulse tone was followed by the 120 db startle tone. In the prepulse.startle trials, the mouse was exposed to the prepulse for 20 ms and the startle for 40 ms with a 100 ms gap between the two tones. Each trial was averaged over the 6 blocks, and percent PPI was calculated as:

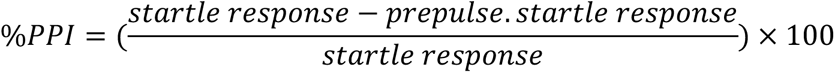

n=8-13 mice/group; age=16-20 weeks

##### Fear Conditioning

To test for deficits in associative learning and memory, we performed a 3-day fear conditioning paradigm. Training and testing were performed in a chamber (H10-11M-TC, Coulbourn Instruments) equipped with a house light, a ceiling-mounted camera, and a speaker. On days 1 and 2, the chamber was equipped with an electric grid shock floor (H10-11M-TC-SF, Coulbourn Instruments); on day 3, the chamber was equipped with a non-shock wire mesh floor (H10-11M-TC-NSF, Coulbourn Instruments). On day 1, the animals were subjected to a 7 min training trial consisting of a 3 min acclimation period followed by three tone-shock pairings during which a tone was played for 20 s immediately followed by a 1 s, 0.5 mA foot shock. On day 2, the animals were placed back in the same chamber as day 1 and were left for 7 min without presentation of the tone or foot shock to test contextual memory. On day 3, the animals were placed in a different chamber, and the chamber floor was replaced with the non-shock wire mesh floor. The animals were in the chamber for 7 min, and the shock-associated tone was played for the last 320 s of the trial to test cued memory. Freezing behavior was automatically recorded using FreezeFrame (Coulbourn Instruments) during each trial. n=9-13 mice/group; age=16-20 weeks

##### Amphetamine-Induced Locomotor Activity

To evaluate amphetamine sensitivity, amphetamine-induced locomotor activity was measured. The assay was performed in a locomotor chamber (San Diego Instruments) consisting of a plexiglass cage (48×25×22 cm) containing corncob bedding. The locomotor chamber was placed inside an apparatus that projected an 8×4 grid of infrared beams, with beams placed 5 cm apart. When a mouse crossed two consecutive beams, it was considered one ambulation. After a 2 hr acclimation period, mice were given an intraperitoneal injection of either saline or 7.5 mg/kg D-amphetamine and post-injection ambulations were recorded for 2 hr with the accompanying Photobeam Activity System software (San Diego Instruments). Treatments were spread over 2 weeks and were randomized, so that not all of the mice received the same injection in a given week. n=9-13 mice/group; age=16-20 weeks

#### Meta-analysis

A PubMed search using the keywords “mouse metabolomics” was conducted on July 7, 2020. Papers were filtered to only include publicly available studies published in English between 2015 and 2020, resulting in a list of 2601 papers. 500 papers were randomly selected using a random number generator for analysis. If a study did not use metabolomics or if a study did not use metabolomics performed on primary mouse tissue or cultured mouse cells, or if the study was not a primary research paper, it was excluded and replaced with another randomly selected study from the list of 2601 papers. Papers were reviewed and coded as one of the following to indicate how sex as a biological variable was addressed: “males only”, “females only”, “sex not specified”, “both sexes, not stratified”, or “stratified”.

### Quantification and Statistical Analysis

Males and females were analyzed separately in all analyses. All data is represented as mean ± standard error of the mean (SEM), and sample size is included in the figure legend. Values of p<0.05 were considered statistically significant. WT was set as the reference genotype and the STD was set as the reference diet for all analyses. All plots were created using the plotly R package (Sievert et al., 2017) unless otherwise specified. All analyses performed in R utilized R 3.5.3 (R Core Team, 2008). All analyses performed in Prism used Prism 8.3.1 (GraphPad).

Specific details for each analysis are as follows:

#### Growth Curves

Growth curve data were analyzed in R (R Core Team, 2008) using the geepack package to implement generalized estimating equations (GEE) that regressed weight measurements on genotype and age while accounting for within-subject correlation of measurements resulting from multiple time points of data collection (Halekoh et al., 2006; Yan, 2002; Yan and Fine, 2004). Age was dichotomized to “early” (1-3 weeks of age) and “late” (4-16 weeks of age) to coincide with time of weaning. All analyses were repeated after applying an inverse normal transformation to the weight data to better satisfy modeling assumptions. Results using the raw and transformed data led to identical conclusions, so the results from the analysis of the raw weight data were presented for ease of interpretation. Using the GEE framework, we performed four distinct sets of analyses. We first compared weight measurements between B6.Del16^+/*Bdh1-Tfrc*^ and WT mice on the STD. We then compared weight measurements between an independent set of B6.Del16^+/*Bdh1-Tfrc*^ and WT mice on the HFD. To test for sex-specific differences in effect size in each diet condition, we pooled the male and female data from that diet treatment and fit an additional GEE model that regressed weight measurements on genotype, age, sex, and a genotype-by-sex interaction term. The interaction term was tested to determine whether the effect size of the B6.Del16^+/*Bdh1-Tfrc*^ genotype significantly differed by sex. Finally, to test for the effect of the HFD intervention on effect size within each sex, we pooled the STD and HFD data from each sex and fit a new GEE model that regressed weight measurements on genotype, age, diet, and a genotype-by-diet interaction term. The interaction term was tested to determine whether the effect size of the B6.Del16^+/*Bdh1-Tfrc*^ genotype significantly differed between diet treatments.

#### Indirect Calorimetry

All data were analyzed in R (R Core Team, 2008). The data were filtered to exclude observations with a respiratory exchange ratio (RER) less than 0.650 or greater than 1.05, because they are outside the dynamic range of the measurement. Observations with a negative value for cumulative food or water consumption were excluded, because they indicate periods where the animal had climbed onto the sensor. The final dataset was trimmed to remove intervals at the beginning and end of the experiment that did not have observations for every subject. Ambulations were analyzed with a simple linear model implemented through the stats R package regressing ambulations on genotype and age (R Core Team, 2008). Ambulations were averaged over all observations in the light and dark cycles for each animal, and light and dark cycle data were analyzed separately. For food and water consumption, the final value for cumulative consumption was used, and was divided by the animal’s body weight to find the g food consumed per g body weight. The relative food and water consumption data were analyzed with simple linear models implemented through the stats R package regressing relative food or water consumption on genotype and age (R Core Team, 2008). For food consumption, a mediation analysis was performed using the R package mediation to determine if body weight mediated the relationship between genotype and total food consumption (Tingley et al., 2014). For energy expenditure and RER, the data were subsetted to only the peaks and troughs in the RER curve; the peak and trough were identified via manual inspection, and one interval to either side of the peak or trough was included, for a total of 3 data points per interval and 7 total intervals (the day 1 and day 5 light cycles were excluded because the entire cycle was not captured). Each interval was analyzed with a GEE implemented through the geepack R package that regressed either energy expenditure or RER on genotype and interval while accounting for within-subject correlation of measurements resulting from multiple time points of data collection (Halekoh et al., 2006; Yan, 2002; Yan and Fine, 2004).

#### Metabolomics

All data were analyzed in R (R Core Team, 2008). Median summarized ComBat batch-corrected data was used for all analyses (Johnson et al., 2007). To determine the similarity between blind replicate samples, correlation tests were performed using the stats R package (R Core Team, 2008). Data filtering and feature selection with partial least squares regression and linear regression were performed using the xmsPANDA R package (Uppal, 2018). Features missing from more than 50% of all samples, or more than 80% of samples from one group, were removed from the dataset. Stepwise feature selection was performed, where the data were first filtered based on a VIP score>1.5, and then filtered again based on a p value<0.05. All features with a p value<0.05 also had a VIP score>1.5, so the p values from linear regression were used for pathway enrichment analysis. Pathway enrichment analysis was performed using mummichog 2.3.3 with a p value<0.05 cutoff (Li et al., 2013). For hierarchical clustering, the linear regression results from the HILIC and C18 columns were pooled, and the top 250 ranked features across the two datasets based on VIP score and p value were used as input. Hierarchical clustering was implemented via the xmsPANDA R package using Spearman correlation (Uppal, 2018). For predictive modeling, we used the R package randomForest (Liaw and Wiener, 2002) and the top 250 ranked metabolic features to train a random forest with 25,001 trees to predict genotype. Trees were grown to the maximum size possible; by default, 15 features were considered as candidates at each split, and splitter importance was calculated as the mean decrease in Gini impurity. To account for unequal sample sizes in the STD data, we used weights equal to the inverse of the sample size for each genotype. The receiver operating characteristic (ROC) curves and the area under the ROC curves (AUC) were generated using the R package ROCR (Sing et al., 2005). P values for the AUC and prediction accuracy were calculated by permuting genotype 10,000 times. Venn diagrams were constructed to compare male and female datasets within a diet condition, as well as to compare STD and HFD datasets within sex, using the VennDiagram R package (Chen, 2018). Male-female comparisons within a diet condition were performed on the top 250 ranked features. STD-HFD comparisons within sex were performed on statistically significant high-confidence annotated features, defined as an xMSannotator confidence level of 2 or 3 (Uppal et al., 2017).

#### Behavior Tests

##### MWM

Animals that did not swim during the probe trial were removed from all analyses. Training data (swim distance, latency to platform, and swim speed) from the STD and HFD cohorts were analyzed separately in Prism (GraphPad) using two-way repeated measures ANOVA followed by multiple comparisons with Sidak’s correction when a significant genotype effect or interaction was observed. Unpaired t-tests were implemented using the stats R package to separately analyze probe trial data (proportion of time the animals spent in the quadrant of the maze that formerly contained the platform) from the STD and HFD cohorts (R Core Team, 2008). To test for the effect of the diet intervention in the training phase, linear mixed-effects models were implemented using the lme4 R package (Bates et al., 2014). When analyzing the training phase data, we fit models with genotype as the predictor and diet and day as covariates, with subject ID as a random effect. We started with a model including up to a three-way interaction between genotype, diet, and day. To identify the most parsimonious model, we performed backward elimination via likelihood ratio tests implemented using the lmtest R package (Zeileis and Hothorn, 2002) and removed any higher-order interaction terms that were not significant and refit the model. We performed this process with three-way interactions followed by two-way interactions if the three-way interaction was not significant. The final models were fit with both maximum likelihood estimation and restricted maximum likelihood estimation; the fits were comparable, so the results from the models fitted with maximum likelihood are presented. P values were calculated using Satterthwaite’s method via the lmerTest R package (Kuznetsova et al., 2017). To test for the effect of the diet intervention in the probe trial, simple linear models regressing the proportion of time spent in the platform quadrant on genotype, diet, and a genotype-by-diet interaction were implemented using the stats R package (R Core Team, 2008).

##### Acoustic Startle

Startle response to 70 db was excluded from the dataset for all animals. The inverse normal transformation was applied to transform the data to an approximately normal distribution. Proper transformation of the data was confirmed with the Shapiro-Wilk normality test implemented using the stats R package (R Core Team, 2008). Linear mixed-effects models were implemented using the lme4 R package (Bates et al., 2014). When analyzing the data for each diet separately, all models included genotype, decibel level, and weight as fixed effects and subject ID as a random effect. The models were fit with both maximum likelihood estimation and restricted maximum likelihood estimation; the fits were comparable, so the results from the models fitted with maximum likelihood are presented. To test for the effect of the diet intervention, we fit models with genotype as the predictor and diet, decibel level, and weight as covariates, with subject ID as a random effect. We started with a model including up to a four-way interaction between genotype, diet, decibel, and weight. To identify the most parsimonious model, we performed backward elimination via likelihood ratio tests implemented using the lmtest R package (Zeileis and Hothorn, 2002) and removed any higher-order interaction terms that were not significant and refit the model. We performed this process with four-way interactions, followed by three-way interactions if the four-way interaction was not significant, and followed by two-way interactions if the three-way interactions were not significant. The final models were fit with both maximum likelihood estimation and restricted maximum likelihood estimation; the fits were comparable, so the results from the models fitted with maximum likelihood are presented. P values were calculated using Satterthwaite’s method via the lmerTest R package (Kuznetsova et al., 2017).

##### PPI

Only the prepulse.startle trials were used to calculate % PPI, as shown in the equation above. The response to the PP1.startle condition (70 db prepulse) was excluded from analysis. Data from the STD and HFD cohorts were analyzed separately in Prism (GraphPad) using 2-way repeated measures ANOVA followed by multiple comparisons with Sidak’s correction when a significant genotype effect or interaction was observed. To test for the effect of the diet intervention, linear mixed-effects models were implemented using the lme4 R package (Bates et al., 2014). We fit models with genotype as the predictor and diet and prepulse decibel as covariates, with subject ID as a random effect. We started with a model including up to a three-way interaction between genotype, diet, and prepulse decibel. To identify the most parsimonious model, we performed backward elimination via likelihood ratio tests implemented using the lmtest R package (Zeileis and Hothorn, 2002) and removed any higher-order interaction terms that were not significant and refit the model. We performed this process with three-way interactions followed by two-way interactions if the three-way interaction was not significant.

The final models were fit with both maximum likelihood estimation and restricted maximum likelihood estimation; the fits were comparable, so the results from the models fitted with maximum likelihood are presented. P values were calculated using Satterthwaite’s method via the lmerTest R package (Kuznetsova et al., 2017).

##### Fear Conditioning

Data from each day of the task were analyzed separately. Data from the STD and HFD cohorts were analyzed separately in Prism (GraphPad) using 2-way repeated measures ANOVA followed by multiple comparisons with Sidak’s correction when a significant genotype effect or interaction was observed. To test for the effect of the diet intervention, linear mixed-effects models were implemented using the lme4 R package (Bates et al., 2014). All models included genotype, diet, and a genotype-by-diet interaction as fixed effects and subject ID and time as random effects. The models were fit with both maximum likelihood estimation and restricted maximum likelihood estimation; the fits were comparable, so the results from the models fitted with maximum likelihood are presented. P values were calculated using Satterthwaite’s method via the lmerTest R package (Kuznetsova et al., 2017).

##### Amphetamine-Induced Locomotor Activity

Because the ambulation data were not normally distributed, the inverse normal function was used to transform the data to an approximately normal distribution. Proper transformation of the data as confirmed with the Shapiro-Wilk normality test implemented using the stats R package (R Core Team, 2008) Linear mixed-effects models were implemented using the lme4 R package (Bates et al., 2014). Saline was set as the reference treatment for all analyses. When analyzing the data for each diet separately, all models included genotype, treatment, and a genotype-by-treatment interaction as fixed effects and subject ID and timepoint as random effects. The models were fit with both maximum likelihood estimation and restricted maximum likelihood estimation; the fits were comparable, so the results from the models fitted with maximum likelihood are presented. To test for the effect of the diet intervention, as well as for differences in the effect of the B6.Del16^+/*Bdh1-Tfrc*^ genotype after the HFD intervention, we fit models with genotype as the predictor and diet and treatment as covariates, with subject ID and time as random effects. We started with a model including up to a three-way interaction between genotype, diet, and treatment. To identify the most parsimonious model, we performed backward elimination via likelihood ratio tests implemented using the lmtest R package (Zeileis and Hothorn, 2002) and removed any higher-order interaction terms that were not significant and refit the model. We performed this process with three-way interactions followed by two-way interactions if the three-way interaction was not significant. The final models were fit with both maximum likelihood estimation and restricted maximum likelihood estimation; the fits were comparable, so the results from the models fitted with maximum likelihood are presented. P values were calculated using Satterthwaite’s method via the lmerTest R package (Kuznetsova et al., 2017).

##### Brain Weight

Data were analyzed in R (R Core Team, 2008). To calculate the brain:body weight ratio, the brain weight was divided by the body weight of the animal at euthanasia. Brain weight and the brain:body weight ratio were analyzed by unpaired t-test using the stats R package (R Core Team, 2008).

## REFERENCES

Abel, K.M., Drake, R., and Goldstein, J.M. (2010). Sex differences in schizophrenia. Int Rev Psychiatry 22, 417–428.

Addis, T., Poo, L.J., and Lew, W. (1936). The quantities of protein lost by the various organs and tissues of the body during a fast. Journal of Biological Chemistry 115, 111–116.

Akil, M., and Brewer, G.J. (1995). Psychiatric and behavioral abnormalities in Wilson’s disease. Adv Neurol 65, 171–178.

Andreazza, A.C., Wang, J.F., Salmasi, F., Shao, L., and Young, L.T. (2013). Specific subcellular changes in oxidative stress in prefrontal cortex from patients with bipolar disorder. J Neurochem 127, 552–561.

Arbogast, T., Ouagazzal, A.M., Chevalier, C., Kopanitsa, M., Afinowi, N., Migliavacca, E., Cowling, B.S., Birling, M.C., Champy, M.F., Reymond, A., et al. (2016). Reciprocal Effects on Neurocognitive and Metabolic Phenotypes in Mouse Models of 16p11.2 Deletion and Duplication Syndromes. PLoS Genet 12, e1005709.

Baba, M., Yokoyama, K., Seiriki, K., Naka, Y., Matsumura, K., Kondo, M., Yamamoto, K., Hayashida, M., Kasai, A., Ago, Y., et al. (2019). Psychiatric-disorder-related behavioral phenotypes and cortical hyperactivity in a mouse model of 3q29 deletion syndrome. Neuropsychopharmacology.

Ballif, B.C., Theisen, A., Coppinger, J., Gowans, G.C., Hersh, J.H., Madan-Khetarpal, S., Schmidt, K.R., Tervo, R., Escobar, L.F., Friedrich, C.A., et al. (2008). Expanding the clinical phenotype of the 3q29 microdeletion syndrome and characterization of the reciprocal microduplication. Molecular Cytogenetics 1, 8.

Bates, D., Mächler, M., Bolker, B., and Walker, S. (2014). Fitting linear mixed-effects models using lme4. arXiv preprint arXiv:1406.5823.

Ben-Shachar, D. (2002). Mitochondrial dysfunction in schizophrenia: a possible linkage to dopamine. J Neurochem 83, 1241–1251.

Ben-Shachar, D., and Karry, R. (2008). Neuroanatomical pattern of mitochondrial complex I pathology varies between schizophrenia, bipolar disorder and major depression. PLoS One 3, e3676.

Ben-Shachar, D., and Laifenfeld, D. (2004). Mitochondria, synaptic plasticity, and schizophrenia. Int Rev Neurobiol 59, 273–296.

Ben-Shachar, D., Zuk, R., and Glinka, Y. (1995). Dopamine neurotoxicity: inhibition of mitochondrial respiration. J Neurochem 64, 718–723.

Bergman, O., and Ben-Shachar, D. (2016). Mitochondrial Oxidative Phosphorylation System (OXPHOS) Deficits in Schizophrenia: Possible Interactions with Cellular Processes. Can J Psychiatry 61, 457–469.

Bilder, D.A., Noel, J.K., Baker, E.R., Irish, W., Chen, Y., Merilainen, M.J., Prasad, S., and Winslow, B.J. (2016). Systematic Review and Meta-Analysis of Neuropsychiatric Symptoms and Executive Functioning in Adults With Phenylketonuria. Dev Neuropsychol 41, 245–260.

Blaak, E. (2001). Gender differences in fat metabolism. Curr Opin Clin Nutr Metab Care 4, 499–502.

Bochukova, E.G., Huang, N., Keogh, J., Henning, E., Purmann, C., Blaszczyk, K., Saeed, S., Hamilton-Shield, J., Clayton-Smith, J., O’Rahilly, S., et al. (2010). Large, rare chromosomal deletions associated with severe early-onset obesity. Nature 463, 666–670.

Brennand, K., Savas, J.N., Kim, Y., Tran, N., Simone, A., Hashimoto-Torii, K., Beaumont, K.G., Kim, H.J., Topol, A., Ladran, I., et al. (2015). Phenotypic differences in hiPSC NPCs derived from patients with schizophrenia. Mol Psychiatry 20, 361–368.

Brunetti-Pierri, N., Berg, J.S., Scaglia, F., Belmont, J., Bacino, C.A., Sahoo, T., Lalani, S.R., Graham, B., Lee, B., Shinawi, M., et al. (2008). Recurrent reciprocal 1q21.1 deletions and duplications associated with microcephaly or macrocephaly and developmental and behavioral abnormalities. Nature Genetics 40, 1466–1471.

Burns, B., Schmidt, K., Williams, S.R., Kim, S., Girirajan, S., and Elsea, S.H. (2010). Rai1 haploinsufficiency causes reduced Bdnf expression resulting in hyperphagia, obesity and altered fat distribution in mice and humans with no evidence of metabolic syndrome. Hum Mol Genet 19, 4026–4042.

Chen, H. (2018). VennDiagram: generate high-resolution venn and euler plots. R package version 1.6.20.

Chen, X., McClusky, R., Chen, J., Beaven, S.W., Tontonoz, P., Arnold, A.P., and Reue, K. (2012). The number of x chromosomes causes sex differences in adiposity in mice. PLoS Genet 8, e1002709.

Chen, X., McClusky, R., Itoh, Y., Reue, K., and Arnold, A.P. (2013). X and Y chromosome complement influence adiposity and metabolism in mice. Endocrinology 154, 1092–1104.

Cox, D.M., and Butler, M.G. (2015). A clinical case report and literature review of the 3q29 microdeletion syndrome. Clinical dysmorphology 24, 89–94.

Cox, D.W. (1999). Disorders of copper transport. Br Med Bull 55, 544–555.

D’Angelo, D., Lebon, S., Chen, Q., Martin-Brevet, S., Snyder, L.G., Hippolyte, L., Hanson, E., Maillard, A.M., Faucett, W.A., Mace, A., et al. (2016). Defining the Effect of the 16p11.2 Duplication on Cognition, Behavior, and Medical Comorbidities. JAMA Psychiatry 73, 20–30.

De Vries, G.J., Rissman, E.F., Simerly, R.B., Yang, L.Y., Scordalakes, E.M., Auger, C.J., Swain, A., Lovell-Badge, R., Burgoyne, P.S., and Arnold, A.P. (2002). A model system for study of sex chromosome effects on sexually dimorphic neural and behavioral traits. J Neurosci 22, 9005–9014.

Doyle, C.M., Channon, S., Orlowska, D., and Lee, P.J. (2010). The neuropsychological profile of galactosaemia. J Inherit Metab Dis 33, 603–609.

Edelman, E.A., Girirajan, S., Finucane, B., Patel, P.I., Lupski, J.R., Smith, A.C., and Elsea, S.H. (2007). Gender, genotype, and phenotype differences in Smith-Magenis syndrome: a meta-analysis of 105 cases. Clin Genet 71, 540–550.

Faulds, M.H., Zhao, C., Dahlman-Wright, K., and Gustafsson, J. (2012). The diversity of sex steroid action: regulation of metabolism by estrogen signaling. J Endocrinol 212, 3–12.

Fernandez, A., Meechan, D.W., Karpinski, B.A., Paronett, E.M., Bryan, C.A., Rutz, H.L., Radin, E.A., Lubin, N., Bonner, E.R., Popratiloff, A., et al. (2019). Mitochondrial Dysfunction Leads to Cortical Under-Connectivity and Cognitive Impairment. Neuron 102, 1127–1142.e1123.

Frye, R.E., Cox, D., Slattery, J., Tippett, M., Kahler, S., Granpeesheh, D., Damle, S., Legido, A., and Goldenthal, M.J. (2016). Mitochondrial Dysfunction may explain symptom variation in Phelan-McDermid Syndrome. Sci Rep 6, 19544.

Frye, R.E., Delatorre, R., Taylor, H., Slattery, J., Melnyk, S., Chowdhury, N., and James, S.J. (2013). Redox metabolism abnormalities in autistic children associated with mitochondrial disease. Transl Psychiatry 3, e273.

Giardino, G., Cirillo, E., Maio, F., Gallo, V., Esposito, T., Naddei, R., Grasso, F., and Pignata, C. (2014). Gastrointestinal involvement in patients affected with 22q11.2 deletion syndrome. Scand J Gastroenterol 49, 274–279.

Glassford, M.R., Rosenfeld, J.A., Freedman, A.A., Zwick, M.E., Mulle, J.G., and Unique Rare Chromosome Disorder Support, G. (2016). Novel features of 3q29 deletion syndrome: Results from the 3q29 registry. American Journal of Medical Genetics. Part A 170A, 999–1006.

Go, Y.M., Walker, D.I., Liang, Y., Uppal, K., Soltow, Q.A., Tran, V., Strobel, F., Quyyumi, A.A., Ziegler, T.R., Pennell, K.D., et al. (2015). Reference Standardization for Mass Spectrometry and High-resolution Metabolomics Applications to Exposome Research. Toxicol Sci 148, 531–543.

Gokhale, A., Hartwig, C., Freeman, A.A.H., Bassell, J.L., Zlatic, S.A., Sapp Savas, C., Vadlamudi, T., Abudulai, F., Pham, T.T., Crocker, A., et al. (2019). Systems Analysis of the 22q11.2 Microdeletion Syndrome Converges on a Mitochondrial Interactome Necessary for Synapse Function and Behavior. J Neurosci 39, 3561–3581.

Halekoh, U., Højsgaard, S., and Yan, J. (2006). The R package geepack for generalized estimating equations. Journal of Statistical Software 15, 1–11.

Hanson, E., Bernier, R., Porche, K., Jackson, F.I., Goin-Kochel, R.P., Snyder, L.G., Snow, A.V., Wallace, A.S., Campe, K.L., Zhang, Y., et al. (2015). The cognitive and behavioral phenotype of the 16p11.2 deletion in a clinically ascertained population. Biological Psychiatry 77, 785–793.

Imrie, J., Dasgupta, S., Besley, G.T., Harris, C., Heptinstall, L., Knight, S., Vanier, M.T., Fensom, A.H., Ward, C., Jacklin, E., et al. (2007). The natural history of Niemann-Pick disease type C in the UK. J Inherit Metab Dis 30, 51–59.

Imrie, J., Vijayaraghaven, S., Whitehouse, C., Harris, S., Heptinstall, L., Church, H., Cooper, A., Besley, G.T., and Wraith, J.E. (2002). Niemann-Pick disease type C in adults. J Inherit Metab Dis 25, 491–500.

Itsara, A., Cooper, G.M., Baker, C., Girirajan, S., Li, J., Absher, D., Krauss, R.M., Myers, R.M., Ridker, P.M., Chasman, D.I., et al. (2009). Population analysis of large copy number variants and hotspots of human genetic disease. American Journal of Human Genetics 84, 148–161.

Jackson, C.M. (1915). Effects of acute and chronic inanition upon the relative weights of the various organs and systems of adult albino rats. The American Journal of Anatomy 18, 75–116.

Jacquemont, S., Reymond, A., Zufferey, F., Harewood, L., Walters, R.G., Kutalik, Z., Martinet, D., Shen, Y., Valsesia, A., Beckmann, N.D., et al. (2011). Mirror extreme BMI phenotypes associated with gene dosage at the chromosome 16p11.2 locus. Nature 478, 97–102.

Johnson, W.E., Li, C., and Rabinovic, A. (2007). Adjusting batch effects in microarray expression data using empirical Bayes methods. Biostatistics 8, 118–127.

Jones, D.P., and Sies, H. (2015). The Redox Code. Antioxid Redox Signal 23, 734–746.

Jones, D.P., Walker, D.I., Uppal, K., Rohrbeck, P., Mallon, C.T., and Go, Y.M. (2016). Metabolic Pathways and Networks Associated With Tobacco Use in Military Personnel. J Occup Environ Med 58, S111–116.

Kamitaki, N., Sekar, A., Handsaker, R.E., de Rivera, H., Tooley, K., Morris, D.L., Taylor, K.E., Whelan, C.W., Tombleson, P., Loohuis, L.M.O., et al. (2020). Complement genes contribute sex-biased vulnerability in diverse disorders. Nature.

Karry, R., Klein, E., and Ben Shachar, D. (2004). Mitochondrial complex I subunits expression is altered in schizophrenia: a postmortem study. Biol Psychiatry 55, 676–684.

Kendall, K.M., Rees, E., Escott-Price, V., Einon, M., Thomas, R., Hewitt, J., O’Donovan, M.C., Owen, M.J., Walters, J.T.R., and Kirov, G. (2017). Cognitive Performance Among Carriers of Pathogenic Copy Number Variants: Analysis of 152,000 UK Biobank Subjects. Biol Psychiatry 82, 103–110.

Keys, A., Brožek, J., Henschel, A., Mickelsen, O., and Taylor, H.L. (1950). The biology of human starvation. (2 vols). (Oxford, England: Univ. of Minnesota Press).

Khramtsova, E.A., Davis, L.K., and Stranger, B.E. (2019). The role of sex in the genomics of human complex traits. Nat Rev Genet 20, 173–190.

Kirov, G., Pocklington, A.J., Holmans, P., Ivanov, D., Ikeda, M., Ruderfer, D., Moran, J., Chambert, K., Toncheva, D., Georgieva, L., et al. (2012). De novo CNV analysis implicates specific abnormalities of postsynaptic signalling complexes in the pathogenesis of schizophrenia. Mol Psychiatry 17, 142–153.

Korner, M., Kalin, S., Zweifel-Zehnder, A., Fankhauser, N., Nuoffer, J.M., and Gautschi, M. (2019). Deficits of facial emotion recognition and visual information processing in adult patients with classical galactosemia. Orphanet J Rare Dis 14, 56.

Kuiper, A., Grunewald, S., Murphy, E., Coenen, M.A., Eggink, H., Zutt, R., Rubio-Gozalbo, M.E., Bosch, A.M., Williams, M., Derks, T.G.J., et al. (2019). Movement disorders and nonmotor neuropsychological symptoms in children and adults with classical galactosemia. J Inherit Metab Dis 42, 451–458.

Kuznetsova, A., Brockhoff, P.B., and Christensen, R.H.B. (2017). lmerTest package: tests in linear mixed effects models. Journal of statistical software 82.

Lacaria, M., Saha, P., Potocki, L., Bi, W., Yan, J., Girirajan, S., Burns, B., Elsea, S., Walz, K., Chan, L., et al. (2012). A duplication CNV that conveys traits reciprocal to metabolic syndrome and protects against diet-induced obesity in mice and men. PLoS Genet 8, e1002713.

Li, S., Park, Y., Duraisingham, S., Strobel, F.H., Khan, N., Soltow, Q.A., Jones, D.P., and Pulendran, B. (2013). Predicting network activity from high throughput metabolomics. PLoS Computational Biology 9, e1003123.

Liaw, A., and Wiener, M. (2002). Classification and regression by randomForest. R news 2, 18–22.

Link, J.C., Chen, X., Arnold, A.P., and Reue, K. (2013). Metabolic impact of sex chromosomes. Adipocyte 2, 74–79.

Link, J.C., and Reue, K. (2017). Genetic Basis for Sex Differences in Obesity and Lipid Metabolism. Annu Rev Nutr 37, 225–245.

Mandy, W., Chilvers, R., Chowdhury, U., Salter, G., Seigal, A., and Skuse, D. (2012). Sex differences in autism spectrum disorder: evidence from a large sample of children and adolescents. J Autism Dev Disord 42, 1304–1313.

Marazziti, D., Baroni, S., Picchetti, M., Landi, P., Silvestri, S., Vatteroni, E., and Catena Dell’Osso, M. (2012). Psychiatric disorders and mitochondrial dysfunctions. Eur Rev Med Pharmacol Sci 16, 270–275.

Marshall, C.R., Howrigan, D.P., Merico, D., Thiruvahindrapuram, B., Wu, W., Greer, D.S., Antaki, D., Shetty, A., Holmans, P.A., Pinto, D., et al. (2017). Contribution of copy number variants to schizophrenia from a genome-wide study of 41,321 subjects. Nature Genetics 49, 27–35.

Matthiesen, N.B., Agergaard, P., Henriksen, T.B., Bach, C.C., Gaynor, J.W., Hjortdal, V., and Ostergaard, J.R. (2016). Congenital Heart Defects and Measures of Fetal Growth in Newborns with Down Syndrome or 22q11.2 Deletion Syndrome. J Pediatr 175, 116–122.e114.

Mayes, J.S., and Watson, G.H. (2004). Direct effects of sex steroid hormones on adipose tissues and obesity. Obes Rev 5, 197–216.

McDonald-McGinn, D.M., Sullivan, K.E., Marino, B., Philip, N., Swillen, A., Vorstman, J.A., Zackai, E.H., Emanuel, B.S., Vermeesch, J.R., Morrow, B.E., et al. (2015). 22q11.2 deletion syndrome. Nat Rev Dis Primers 1, 15071.

Meeng, M., Knobbe, A., Koopman, A., Klinken, J., and Van den Berg, S. (2010). Equation Discovery for Whole-Body Metabolism Modelling. Machine Learning in Systems Biology, 35.

Mefford, H.C., Sharp, A.J., Baker, C., Itsara, A., Jiang, Z., Buysse, K., Huang, S., Maloney, V.K., Crolla, J.A., Baralle, D., et al. (2008). Recurrent rearrangements of chromosome 1q21.1 and variable pediatric phenotypes. N Engl J Med 359, 1685–1699.

Mervis, C.B., Klein-Tasman, B.P., Huffman, M.J., Velleman, S.L., Pitts, C.H., Henderson, D.R., Woodruff-Borden, J., Morris, C.A., and Osborne, L.R. (2015). Children with 7q11.23 duplication syndrome: psychological characteristics. American Journal of Medical Genetics. Part A 167, 1436–1450.

Ming, X., Stein, T.P., Barnes, V., Rhodes, N., and Guo, L. (2012). Metabolic perturbance in autism spectrum disorders: a metabolomics study. J Proteome Res 11, 5856–5862.

Moghadasian, M.H. (2004). Cerebrotendinous xanthomatosis: clinical course, genotypes and metabolic backgrounds. Clin Invest Med 27, 42–50.

Morris, G., and Berk, M. (2015). The many roads to mitochondrial dysfunction in neuroimmune and neuropsychiatric disorders. BMC Med 13, 68.

Mulle, J.G. (2015). The 3q29 deletion confers >40-fold increase in risk for schizophrenia. Molecular Psychiatry 20, 1028–1029.

Mulle, J.G., Dodd, A.F., McGrath, J.A., Wolyniec, P.S., Mitchell, A.A., Shetty, A.C., Sobreira, N.L., Valle, D., Rudd, M.K., Satten, G., et al. (2010). Microdeletions of 3q29 confer high risk for schizophrenia. American Journal of Human Genetics 87, 229–236.

Napoli, E., Tassone, F., Wong, S., Angkustsiri, K., Simon, T.J., Song, G., and Giulivi, C. (2015). Mitochondrial Citrate Transporter-dependent Metabolic Signature in the 22q11.2 Deletion Syndrome. J Biol Chem 290, 23240–23253.

Ngo, S.T., Steyn, F.J., and McCombe, P.A. (2014). Gender differences in autoimmune disease. Front Neuroendocrinol 35, 347–369.

Norkett, R., Modi, S., Birsa, N., Atkin, T.A., Ivankovic, D., Pathania, M., Trossbach, S.V., Korth, C., Hirst, W.D., and Kittler, J.T. (2016). DISC1-dependent Regulation of Mitochondrial Dynamics Controls the Morphogenesis of Complex Neuronal Dendrites. J Biol Chem 291, 613–629.

O’Brien, S., Repp, A.C., Williams, G.E., and Christophersen, E.R. (1991). Pediatric feeding disorders. Behavior Modification 15, 394–418.

O’Sullivan, A.J., Crampton, L.J., Freund, J., and Ho, K.K. (1998). The route of estrogen replacement therapy confers divergent effects on substrate oxidation and body composition in postmenopausal women. J Clin Invest 102, 1035–1040.

Ockner, R.K., Burnett, D.A., Lysenko, N., and Manning, J.A. (1979). Sex differences in long chain fatty acid utilization and fatty acid binding protein concentration in rat liver. J Clin Invest 64, 172–181.

Ockner, R.K., Lysenko, N., Manning, J.A., Monroe, S.E., and Burnett, D.A. (1980). Sex steroid modulation of fatty acid utilization and fatty acid binding protein concentration in rat liver. J Clin Invest 65, 1013–1023.

Palmisano, B.T., Zhu, L., Eckel, R.H., and Stafford, J.M. (2018). Sex differences in lipid and lipoprotein metabolism. Mol Metab 15, 45–55.

Park, C., Lee, S.A., Hong, J.H., Suh, Y., Park, S.J., Suh, B.K., Woo, Y., Choi, J., Huh, J.W., Kim, Y.M., et al. (2016). Disrupted-in-schizophrenia 1 (DISC1) and Syntaphilin collaborate to modulate axonal mitochondrial anchoring. Mol Brain 9, 69.

Park, C., and Park, S.K. (2012). Molecular links between mitochondrial dysfunctions and schizophrenia. Mol Cells 33, 105–110.

Patterson, M.C., Hendriksz, C.J., Walterfang, M., Sedel, F., Vanier, M.T., and Wijburg, F. (2012). Recommendations for the diagnosis and management of Niemann-Pick disease type C: an update. Mol Genet Metab 106, 330–344.

Pettersson, M., Viljakainen, H., Loid, P., Mustila, T., Pekkinen, M., Armenio, M., Andersson-Assarsson, J.C., Makitie, O., and Lindstrand, A. (2017). Copy Number Variants Are Enriched in Individuals With Early-Onset Obesity and Highlight Novel Pathogenic Pathways. J Clin Endocrinol Metab 102, 3029–3039.

Pinero-Martos, E., Ortega-Vila, B., Pol-Fuster, J., Cisneros-Barroso, E., Ruiz-Guerra, L., Medina-Dols, A., Heine-Suner, D., Llado, J., Olmos, G., and Vives-Bauza, C. (2016). Disrupted in schizophrenia 1 (DISC1) is a constituent of the mammalian mitochondrial contact site and cristae organizing system (MICOS) complex, and is essential for oxidative phosphorylation. Hum Mol Genet 25, 4157–4169.

Pollak, R.M., Murphy, M.M., Epstein, M.P., Zwick, M.E., Klaiman, C., Saulnier, C.A., and Mulle, J.G. (2019). Neuropsychiatric phenotypes and a distinct constellation of ASD features in 3q29 deletion syndrome: results from the 3q29 registry. Molecular Autism 10, 30.

Purcell, R.H., Sun, B., Pass, L.L., Power, M.L., Moran, T.H., and Tamashiro, K.L. (2011). Maternal stress and high-fat diet effect on maternal behavior, milk composition, and pup ingestive behavior. Physiol Behav 104, 474–479.

R Core Team (2008). R: A language and environment for statistical computing. R Foundation for Statistical Computing, Vienna, Austria.

Rajasekaran, A., Venkatasubramanian, G., Berk, M., and Debnath, M. (2015). Mitochondrial dysfunction in schizophrenia: pathways, mechanisms and implications. Neurosci Biobehav Rev 48, 10–21.

Regitz-Zagrosek, V., and Kararigas, G. (2017). Mechanistic Pathways of Sex Differences in Cardiovascular Disease. Physiol Rev 97, 1–37.

Robicsek, O., Karry, R., Petit, I., Salman-Kesner, N., Muller, F.J., Klein, E., Aberdam, D., and Ben-Shachar, D. (2013). Abnormal neuronal differentiation and mitochondrial dysfunction in hair follicle-derived induced pluripotent stem cells of schizophrenia patients. Mol Psychiatry 18, 1067–1076.

Rosenfeld, M., Brenner-Lavie, H., Ari, S.G., Kavushansky, A., and Ben-Shachar, D. (2011). Perturbation in mitochondrial network dynamics and in complex I dependent cellular respiration in schizophrenia. Biol Psychiatry 69, 980–988.

Rutkowski, T.P., Purcell, R.H., Pollak, R.M., Grewenow, S.M., Gafford, G.M., Malone, T., Khan, U.A., Schroeder, J.P., Epstein, M.P., Bassell, G.J., et al. (2019). Behavioral changes and growth deficits in a CRISPR engineered mouse model of the schizophrenia-associated 3q29 deletion. Molecular Psychiatry.

Sanders, S.J., He, X., Willsey, A.J., Ercan-Sencicek, A.G., Samocha, K.E., Cicek, A.E., Murtha, M.T., Bal, V.H., Bishop, S.L., Dong, S., et al. (2015). Insights into Autism Spectrum Disorder Genomic Architecture and Biology from 71 Risk Loci. Neuron 87, 1215–1233.

Schmidt, C.F., Kety, S.S., and Pennes, H.H. (1945). THE GASEOUS METABOLISM OF THE BRAIN OF THE MONKEY. American Journal of Physiology-Legacy Content 143, 33–52.

Schneider, M., Debbané, M., Bassett, A.S., Chow, E.W.C., Fung, W.L.A., van den Bree, M., Owen, M., Murphy, K.C., Niarchou, M., Kates, W.R., et al. (2014). Psychiatric disorders from childhood to adulthood in 22q11.2 deletion syndrome: results from the International Consortium on Brain and Behavior in 22q11.2 Deletion Syndrome. The American Journal of Psychiatry 171, 627–639.

Sefik, E., Purcell, R.H., Merritt-Garza, M., Karne, S., Randall, J., Walker, E.F., Bassell, G.J., and Mulle, J.G. (2020). Convergent and distributed effects of the schizophrenia-associated 3q29 deletion on the human neural transcriptome. bioRxiv, 2020.2005.2025.111351.

Sievert, C., Parmer, C., Hocking, T., Chamberlain, S., Ram, K., Corvellec, M., and Despouy, P. (2017). plotly: Create Interactive Web Graphics via ‘plotly.js’. R package version 4.6.0.

Silverman, A.H., and Tarbell, S. (2009). Feeding and vomiting problems in pediatric populations. Handbook of pediatric psychology, 429–445.

Sing, T., Sander, O., Beerenwinkel, N., and Lengauer, T. (2005). ROCR: visualizing classifier performance in R. Bioinformatics 21, 3940–3941.

Smith, A.C., Gropman, A.L., Bailey-Wilson, J.E., Goker-Alpan, O., Elsea, S.H., Blancato, J., Lupski, J.R., and Potocki, L. (2002). Hypercholesterolemia in children with Smith-Magenis syndrome: del (17) (p11.2p11.2). Genet Med 4, 118–125.

Soler-Argilaga, C., and Heimberg, M. (1976). Comparison of metabolism of free fatty acid by isolated perfused livers from male and female rats. J Lipid Res 17, 605–615.

Stefansson, H., Meyer-Lindenberg, A., Steinberg, S., Magnusdottir, B., Morgen, K., Arnarsdottir, S., Bjornsdottir, G., Walters, G.B., Jonsdottir, G.A., Doyle, O.M., et al. (2014). CNVs conferring risk of autism or schizophrenia affect cognition in controls. Nature 505, 361–366.

Szatkiewicz, J.P., O’Dushlaine, C., Chen, G., Chambert, K., Moran, J.L., Neale, B.M., Fromer, M., Ruderfer, D., Akterin, S., Bergen, S.E., et al. (2014). Copy number variation in schizophrenia in Sweden. Molecular Psychiatry 19, 762.

Tarlungeanu, D.C., Deliu, E., Dotter, C.P., Kara, M., Janiesch, P.C., Scalise, M., Galluccio, M., Tesulov, M., Morelli, E., Sonmez, F.M., et al. (2016). Impaired Amino Acid Transport at the Blood Brain Barrier Is a Cause of Autism Spectrum Disorder. Cell 167, 1481–1494.e1418.

Tingley, D., Yamamoto, T., Hirose, K., Keele, L., and Imai, K. (2014). Mediation: R package for causal mediation analysis.

Uppal, K. (2018). xmsPANDA: Predictive And Network Discovery Analysis (xmsPANDA): R package for biomarker discovery and biomarker-driven network analysis for studying disease mechanisms using metabolomics. R package version 1.0.7.5.

Uppal, K., Soltow, Q.A., Strobel, F.H., Pittard, W.S., Gernert, K.M., Yu, T., and Jones, D.P. (2013). xMSanalyzer: automated pipeline for improved feature detection and downstream analysis of large-scale, non-targeted metabolomics data. BMC Bioinformatics 14, 15.

Uppal, K., Walker, D.I., and Jones, D.P. (2017). xMSannotator: an R package for network-based annotation of high-resolution metabolomics data. Analytical chemistry 89, 1063–1067.

Vanier, M.T. (2010). Niemann-Pick disease type C. Orphanet J Rare Dis 5, 16.

Volkert, V.M., Patel, M.R., and Peterson, K.M. (2016). Food refusal and selective eating. In Behavioral health promotion and intervention in intellectual and developmental disabilities (Springer), pp. 137–161.

Voll, S.L., Boot, E., Butcher, N.J., Cooper, S., Heung, T., Chow, E.W., Silversides, C.K., and Bassett, A.S. (2017). Obesity in adults with 22q11.2 deletion syndrome. Genet Med 19, 204–208.

Walsh, B.W., Schiff, I., Rosner, B., Greenberg, L., Ravnikar, V., and Sacks, F.M. (1991). Effects of postmenopausal estrogen replacement on the concentrations and metabolism of plasma lipoproteins. N Engl J Med 325, 1196–1204.

Walters, R.G., Jacquemont, S., Valsesia, A., de Smith, A.J., Martinet, D., Andersson, J., Falchi, M., Chen, F., Andrieux, J., Lobbens, S., et al. (2010). A new highly penetrant form of obesity due to deletions on chromosome 16p11.2. Nature 463, 671–675.

Wesseling, H., Xu, B., Want, E.J., Holmes, E., Guest, P.C., Karayiorgou, M., Gogos, J.A., and Bahn, S. (2017). System-based proteomic and metabonomic analysis of the Df(16)A(+/-) mouse identifies potential miR-185 targets and molecular pathway alterations. Mol Psychiatry 22, 384–395.

Willatt, L., Cox, J., Barber, J., Cabanas, E.D., Collins, A., Donnai, D., FitzPatrick, D.R., Maher, E., Martin, H., Parnau, J., et al. (2005). 3q29 microdeletion syndrome: clinical and molecular characterization of a new syndrome. American Journal of Human Genetics 77, 154–160.

Wraith, J.E., Baumgartner, M.R., Bembi, B., Covanis, A., Levade, T., Mengel, E., Pineda, M., Sedel, F., Topcu, M., Vanier, M.T., et al. (2009). Recommendations on the diagnosis and management of Niemann-Pick disease type C. Mol Genet Metab 98, 152–165.

Wu, B.N., and O’Sullivan, A.J. (2011). Sex differences in energy metabolism need to be considered with lifestyle modifications in humans. J Nutr Metab 2011, 391809.

Yan, J. (2002). Geepack: yet another package for generalized estimating equations. R-news 2, 12–14.

Yan, J., and Fine, J. (2004). Estimating equations for association structures. Statistics in medicine 23, 859–874.

Yap, I.K., Angley, M., Veselkov, K.A., Holmes, E., Lindon, J.C., and Nicholson, J.K. (2010). Urinary metabolic phenotyping differentiates children with autism from their unaffected siblings and age-matched controls. J Proteome Res 9, 2996–3004.

Yu, T., Park, Y., Li, S., and Jones, D.P. (2013). Hybrid feature detection and information accumulation using high-resolution LC-MS metabolomics data. Journal of Proteome Research 12, 1419–1427.

Zeileis, A., and Hothorn, T. (2002). Diagnostic checking in regression relationships.

Zhang, Y., Klein, K., Sugathan, A., Nassery, N., Dombkowski, A., Zanger, U.M., and Waxman, D.J. (2011). Transcriptional profiling of human liver identifies sex-biased genes associated with polygenic dyslipidemia and coronary artery disease. PLoS One 6, e23506.

Zufferey, F., Sherr, E.H., Beckmann, N.D., Hanson, E., Maillard, A.M., Hippolyte, L., Macé, A., Ferrari, C., Kutalik, Z., Andrieux, J., et al. (2012). A 600 kb deletion syndrome at 16p11.2 leads to energy imbalance and neuropsychiatric disorders. J Med Genet 49, 660–668.

