## Supplementary material for "Metabolic effects of the schizophrenia-associated 3q29 deletion are sex-specific and uncoupled from behavioral phenotypes": Key Resources Table

| REAGENT or RESOURCE | SOURCE | IDENTIFIER |
| --- | --- | --- |
| Experimental Models: Organisms/Strains | | |
| Mouse: Del16^+/^*^Bdh1-Tfrc^*: C57BL/6N-Del16^+/^*^Bdh1-Tfrc^* | Rutkowski et al., 2019 | MGI:6241487 |
| Software and Algorithms | | |
| R 3.5.3 | R Foundation for Statistical Computing | https://cran.r-project.org |
| R 3.6.2 | R Foundation for Statistical Computing | https://cran.r-project.org |
| Mummichog 2.3.3 | Li et al., 2013 | http://mummichog.org |
| Prism 8.3.1 | GraphPad | http://www.graphpad.com/scientific-software/prism; RRID: SCR_015807 |
| Microsoft Excel (version 16.37) | Microsoft Corp. | https://www.microsoft.com/en-gb/; RRID: SCR_016137 |
| TopScan | CleverSys | http://cleversysinc.com/CleverSysInc/?csi_products=topscan-suite; RRID: SCR_017141 |
| FreezeFrame | Coulbourn Instruments | https://www.coulbourn.com/category_s/277.htm |
| Deposited Data | | |
| Raw data | | http://dx.doi.org/10.17632/g3v2nt657z.1 |
| Other | | |
| Laboratory Rodent Diet | LabDiet | Cat#5001 |
| Teklad Rodent Diet | ENVIGO | Cat#88137 |
| Oxymax CLAMS-HC | Columbus Instruments | http://www.colinst.com/docs/CLAMS-HC-CF-WC-2018.pdf |
| SR-LAB startle response system | San Diego Instruments | https://sandiegoinstruments.com/product/sr-lab-startle-response/ |
| Mouse test cage | Coulbourn Instruments | Cat#H10-11M-TC |
| Shock floor for mouse test cage | Coulbourn Instruments | Cat#H10-11M-TC-SF |
| Non-shock floor for mouse test cage | Coulbourn Instruments | Cat#H10-11M-TC-NSF |
| Photobeam Activity System - Home Cage | San Diego Instruments | https://sandiegoinstruments.com/wp-content/uploads/2018/08/PAS-Home-Cage-DataSheet.pdf |
