## Supplemental Information for "Metabolic effects of the schizophrenia-associated 3q29 deletion are sex-specific and uncoupled from behavioral phenotypes"

**Energy and water consumption are not reduced in B6.Del16^+/^*^Bdh1-Tfrc^* mice on the STD**

We performed 5 days of indirect calorimetry on male and female WT and B6.Del16^+/^*^Bdh1-Tfrc^* mice using CLAMS/Metabolic Cages (Columbus Instruments). To determine whether the weight deficit in B6.Del16^+/^*^Bdh1-Tfrc^* mice is simply due to decreased calorie intake, we evaluated food consumption. There was no significant difference in food consumption (Figure S1A) between male or female WT and B6.Del16^+/^*^Bdh1-Tfrc^* animals after controlling for the reduced weight of B6.Del16^+/^*^Bdh1-Tfrc^* animals; mediation analysis showed that including weight in the model fully accounted for the relationship between genotype and food consumption. Additionally, there were no differences between male or female WT and B6.Del16^+/^*^Bdh1-Tfrc^* mice in water consumption (Figure S1B).

**Figure S1. Indirect calorimetry of WT and B6.Del16^+/^*^Bdh1-Tfrc^* mice on the STD and HFD. Related to Figures 2 and 4.**

A) Total food consumption for male and female WT (n= 11 male, 14 female) and B6.Del16^+/^*^Bdh1-Tfrc^* (n= 8 male, 12 female) mice on the STD over 5 days in CLAMS/Metabolic Cages.

B) Total water consumption for male and female WT and B6.Del16^+/^*^Bdh1-Tfrc^* mice on the STD over 5 days in CLAMS/Metabolic Cages.

C and D) Average ambulations during the light and dark cycles for C) male and D) female WT and B6.Del16^+/^*^Bdh1-Tfrc^* mice on the STD over 5 days in CLAMS/Metabolic Cages.

E) Total food consumption for male and female WT (n= 10 male, 10 female) and B6.Del16^+/^*^Bdh1-Tfrc^* (n= 10 male, 10 female) mice on the HFD over 5 days in CLAMS/Metabolic Cages.

F) Total water consumption for male and female WT and B6.Del16^+/^*^Bdh1-Tfrc^* mice on the HFD over 5 days in CLAMS/Metabolic Cages.

G and H) Average ambulations during the light and dark cycles for G) male and H) female WT and B6.Del16^+/^*^Bdh1-Tfrc^* mice on the HFD over 5 days in CLAMS/Metabolic Cages.

I and J) Energy expenditure for I) male and J) female WT and B6.Del16^+/^*^Bdh1-Tfrc^* mice on the HFD over 5 days in CLAMS/Metabolic Cages.

Data are represented as mean ± SEM. n.s., p>0.05; *, p<0.05

Statistical analysis of food consumption, water consumption, and ambulations (A-H) was performed using simple linear regression. Statistical analyses of energy expenditure (I and J) were performed using generalized linear models.

**Sex-specific and diet-specific small molecule changes identified via untargeted metabolomics**

We performed untargeted metabolomics of liver samples from WT and B6.Del16^+/^*^Bdh1-Tfrc^* mice on the STD and HFD. To understand the effect of sex on the metabolic environment of animals on the STD, we compared all nominally significant features between males and females. Only 22 features were identified in both datasets (Figure 3A). Of those 22 features, 13 were enriched in samples from B6.Del16^+/^*^Bdh1-Tfrc^* animals and 9 were depleted in samples from B6.Del16^+/^*^Bdh1-Tfrc^* animals. An additional 17 features were identified in both datasets but were discordant, so they were not counted as common features. Of the 17 discordant features, 9 were enriched in samples from B6.Del16^+/^*^Bdh1-Tfrc^* males but depleted in samples from B6.Del16^+/^*^Bdh1-Tfrc^* females, while the other 8 were depleted in samples from B6.Del16^+/^*^Bdh1-Tfrc^* males but enriched in samples from B6.Del16^+/^*^Bdh1-Tfrc^* females. When we compared all nominally significant features between samples from HFD-treated males and females, a similar pattern emerged, with only 7 features identified in both datasets (Figure 5A). Of those 7 features, 6 were enriched in samples from B6.Del16^+/^*^Bdh1-Tfrc^* animals and one was depleted in samples from B6.Del16^+/^*^Bdh1-Tfrc^* animals. An additional 18 features were identified in both datasets but were discordant, so they were not counted as common features. Of the 18 discordant features, 7 were enriched in samples from B6.Del16^+/^*^Bdh1-Tfrc^* males but depleted in samples from B6.Del16^+/^*^Bdh1-Tfrc^* females, while the other 11 were depleted in samples from B6.Del16^+/^*^Bdh1-Tfrc^* males but enriched in samples from B6.Del16^+/^*^Bdh1-Tfrc^* females.

To understand the effect of the HFD on the metabolic environment, we compared the statistically significant high-confidence annotated features between STD-treated and HFD-treated males, and between STD-treated and HFD-treated females. One feature was enriched in samples from B6.Del16^+/^*^Bdh1-Tfrc^* animals in both male datasets (Figure 5H). Three additional features were identified in both datasets but were discordant, so they were not counted as common features. Of the three discordant features, two were enriched in samples from STD-treated B6.Del16^+/^*^Bdh1-Tfrc^* males but depleted in samples from HFD-treated B6.Del16^+/^*^Bdh1-Tfrc^* males, and one was depleted in samples from STD-treated B6.Del16^+/^*^Bdh1-Tfrc^* males but enriched in samples from HFD-treated B6.Del16^+/^*^Bdh1-Tfrc^* males. One feature was depleted in samples from B6.Del16^+/^*^Bdh1-Tfrc^* animals in both female datasets (Figure 5G). An additional two features were identified in both datasets but were discordant, so they were not counted as common features. Both discordant features were depleted in samples from STD-treated B6.Del16^+/^*^Bdh1-Tfrc^* females and enriched in samples from HFD-treated B6.Del16^+/^*^Bdh1-Tfrc^* females.

To evaluate the quality of the metabolomics data, we included four blind replicate samples in the STD experiment and two blind replicate samples in the HFD experiment. The STD replicate samples had an R^2^ value between 0.832 and 0.996, with a mean R^2^ of 0.963±0.055 (Figure S2A-H). The HFD replicate samples had an R^2^ value between 0.987 and 0.997, with a mean R^2^ of 0.991±0.005 (Figure S2I-L). These data demonstrate that there was good replication and consistently high data quality at the feature level across both untargeted metabolomics experiments.

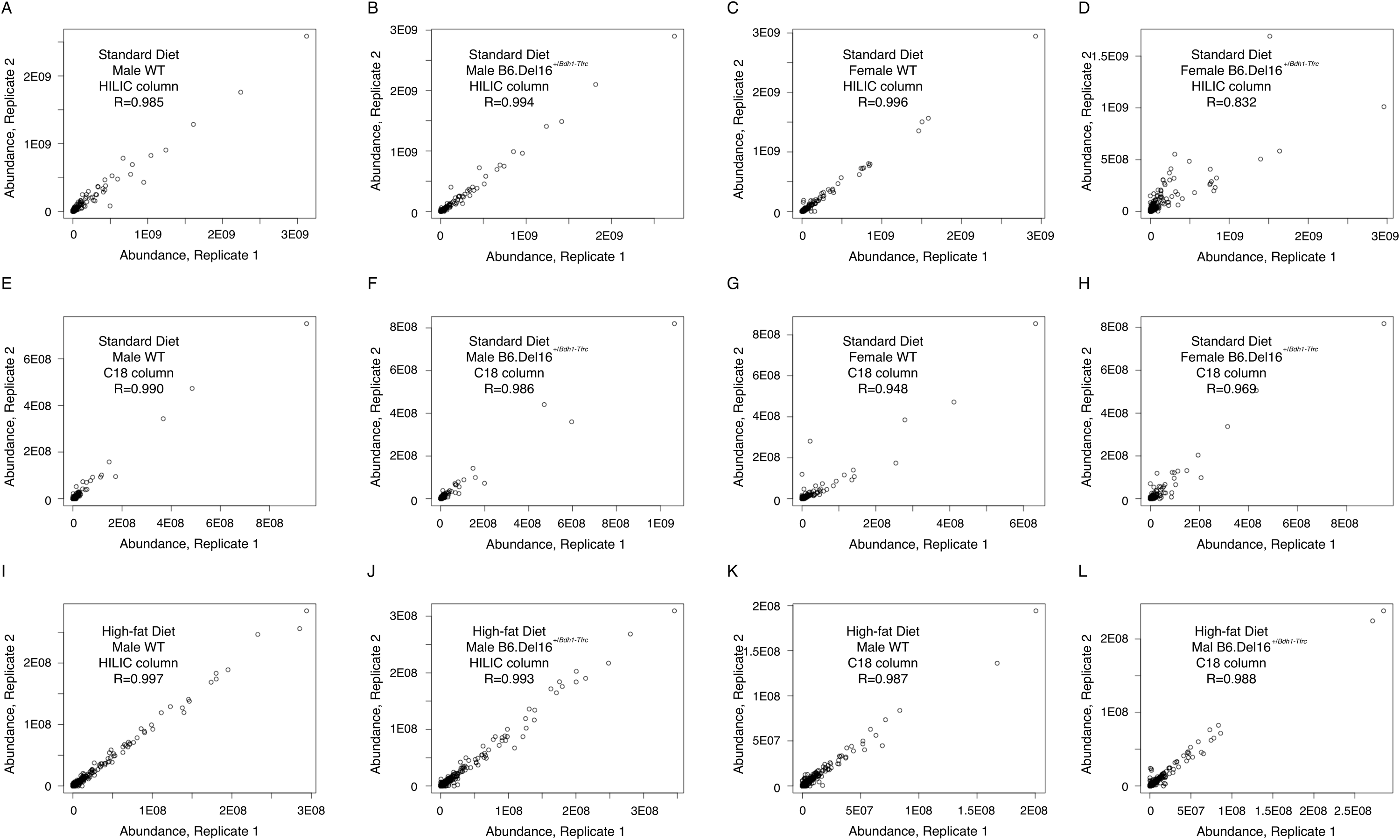

**Figure S2. Correlation between replicate samples from untargeted metabolomics.**

A, B, C, D, E, F, G, and H) Correlation between blind replicate samples for liver metabolomics of STD-treated animals using A, B, C, D) the HILIC column and E, F, G, H) the C18 column.

I, J, K, and L) Correlation between blind replicate samples for liver metabolomics of HFD-treated animals using I, J) the HILIC column and K, L) the C18 column.

**Behavioral phenotypes in B6.Del16^+/^*^Bdh1-Tfrc^* mice are not impacted by HFD treatment**

***Spatial learning and memory***

In the Morris water maze (MWM), we found no differences between male WT and B6.Del16^+/^*^Bdh1-Tfrc^* mice on the STD in swimming distance (p>0.05), latency (p>0.05), or swim speed (p>0.05) during the training portion (Figure S3A-C). Male B6.Del16^+/^*^Bdh1-Tfrc^* mice on the HFD showed increased latency (p=0.002) and swam a greater distance (p=0.002) to reach the hidden platform compared to WT littermates, but did not show any difference in swim speed (p>0.05) during the training portion (Figure S3G-I). When the data from STD- and HFD-treated males were directly compared, we observed a significant main effect of genotype on latency and swim distance (p<0.05), where male B6.Del16^+/^*^Bdh1-Tfrc^* mice took longer to reach the platform and swam a farther distance compared to WT littermates, and a significant main effect of diet on swim distance and swim speed (p<0.05), where males on the HFD swam a shorter distance and swam more slowly than males on the STD (Figure S3A-C, G-I).

Female B6.Del16^+/^*^Bdh1-Tfrc^* mice on the STD showed increased swimming distance (p=0.005), but no differences in latency (p>0.05) or swim speed (p>0.05) compared to WT littermates in the training portion of the MWM (Figure S3D-F). Female B6.Del16^+/^*^Bdh1-Tfrc^* mice on the HFD showed increased latency (p=0.025), but no differences in distance (p>0.05) or swim speed (p>0.05) compared to WT littermates in the training portion of the MWM (Figure S3J-L). When data from STD- and HFD-treated females were directly compared, we observed a significant main effect of genotype on latency and swim distance (p<0.05), where female B6.Del16^+/^*^Bdh1-Tfrc^* mice took longer to reach the platform and swam a farther distance compared to WT littermates, and a significant main effect of diet on latency, where females on the HFD took longer to reach the platform than females on the STD (Figure S3D-F, J-L).

In the probe trial of the MWM, there was no difference in the percentage of time in the quadrant that formerly contained the platform between male or female WT and B6.Del16^+/^*^Bdh1-Tfrc^* mice on the STD (Figure S3M-N). There was no difference in the percentage of time in the quadrant that formerly contained the platform between male B6.Del16^+/^*^Bdh1-Tfrc^* and WT mice on the HFD (p>0.05); however, female B6.Del16^+/^*^Bdh1-Tfrc^* mice on the HFD spent significantly less time in the platform quadrant compared to WT littermates (p=0.022, Figure S3M-N). When the data from STD- and HFD-treated males were directly compared, we found a significant main effect of diet, where males on the HFD spent more time in the quadrant that formerly contained the platform than males on the STD (p=0.02). When the data from STD- and HFD-treated females were directly compared, there were no main effects of genotype or diet (p>0.05, Figure S3M-N).

***Contextual learning and memory***

In the fear conditioning assay, male B6.Del16^+/^*^Bdh1-Tfrc^* mice on the STD showed no significant differences in freezing percentage in the training phase, but showed decreased freezing compared to WT littermates in both the context (p=0.01) and cue (p=0.004) phases (Figure S3O-Q). Likewise, male B6.Del16^+/^*^Bdh1-Tfrc^* mice on the HFD showed no significant differences in freezing percentage in the training phase, but showed decreased freezing compared to WT littermates in both the context (p=0.001) and cue (p=0.006) phases (Figure S3U-W). When the data from STD- and HFD-treated males were directly compared, we found no main effects of genotype or diet in the training phase (p>0.05, Figure S3O, U). In the context phase, there was a significant main effect of genotype, where male B6.Del16^+/^*^Bdh1-Tfrc^* mice showed decreased freezing compared to WT littermates (p=0.002, Figure S3P, V). In the tone phase, there was a significant main effect of genotype, where male B6.Del16^+/^*^Bdh1-Tfrc^* mice showed decreased freezing compared to WT littermates (p=0.001), and a significant main effect of diet, where male animals on the HFD showed increased freezing relative to males on the STD (p=0.008, Figure S3Q, W).

Female B6.Del16^+/^*^Bdh1-Tfrc^* mice on the STD had similar freezing percentages to WT littermates during all phases of the fear conditioning assay (p>0.05, Figure S3R-T). Likewise, female B6.Del16^+/^*^Bdh1-Tfrc^* mice on the HFD had similar freezing percentages to WT littermates during all phases of the task (p>0.05, Figure S3X-Z). When the data from STD- and HFD-treated females were directly compared, we found no main effects of genotype or diet in the training phase (p>0.05, Figure S3R, X). In the context phase there were no main effects of genotype or diet (p>0.05, Figure S3S, Y). In the tone phase there was a significant main effect of diet, where females on the HFD showed increased freezing relative to females on the STD (p=0.007, Figure S3T, Z).

***Acoustic startle response and sensorimotor gating***

Male and female B6.Del16^+/^*^Bdh1-Tfrc^* mice on the STD showed an increased acoustic startle response compared to WT littermates (p<0.05, FigureS4A, C). Body weight was not significantly associated with startle response for male or female animals on the STD (p>0.05), indicating that the reduced weight phenotype in B6.Del16^+/^*^Bdh1-Tfrc^* animals on the STD did not affect the measured startle response. Male and female B6.Del16^+/^*^Bdh1-Tfrc^* mice on the HFD showed an increase in acoustic startle response compared to WT littermates (p<0.05, Figure S4B, D). For male animals on the HFD, body weight was not associated with startle response (p>0.05). Weight was significantly associated with startle response in female animals on the HFD (p=0.001), suggesting that the decreased weight in female B6.Del16^+/^*^Bdh1-Tfrc^* mice on the HFD may have impacted the measured startle response. When the data from STD- and HFD-treated males were directly compared, we found a significant main effect of diet, where males on the HFD showed an increased acoustic startle response compared to males on the STD (p=0.02, Figure S4A-B). When the data from STD- and HFD-treated females were directly compared, we found a significant main effect of diet, where females on the HFD showed a decreased acoustic startle response compared to females on the STD (p=0.03), and a significant main effect of genotype, where female B6.Del16^+/^*^Bdh1-Tfrc^* mice showed a decreased acoustic startle response compared to WT littermates (p=0.02, Figure S4C-D).

Male B6.Del16^+/^*^Bdh1-Tfrc^* mice on the STD showed reduced prepulse inhibition (PPI) compared to WT littermates (p=0.02, Figure S4E), indicating a mild impairment in sensorimotor gating. Female B6.Del16^+/^*^Bdh1-Tfrc^* mice on the STD showed similar PPI to WT littermates at all prepulse levels (p>0.05, Figure S4G). Likewise, male B6.Del16^+/^*^Bdh1-Tfrc^* mice on the HFD showed significantly reduced PPI compared to WT littermates (p=0.003, Figure S4F), while female B6.Del16^+/^*^Bdh1-Tfrc^* mice on the HFD showed similar PPI to WT littermates (p>0.05, Figure S4H). When the data from STD- and HFD-treated males were directly compared, we found a significant main effect of genotype, where male B6.Del16^+/^*^Bdh1-Tfrc^* mice showed significantly reduced PPI compared to WT littermates (p=1.4E-6, Figure S4E-F). When the data from STD- and HFD-treated females were directly compared, there were no significant effects of genotype or diet (p>0.05, Figure S4G-H).

***Amphetamine sensitivity***

In the amphetamine-induced locomotor activity task, there were no differences in ambulatory activity following saline administration between male B6.Del16^+/^*^Bdh1-Tfrc^* mice on the STD and WT littermates (p>0.05). Likewise, there were no differences in ambulatory activity following saline administration between male B6.Del16^+/^*^Bdh1-Tfrc^* mice on the HFD and WT littermates (p>0.05). After administration of 7.5 mg/kg amphetamine, male B6.Del16^+/^*^Bdh1-Tfrc^* mice on the STD showed similar levels of amphetamine-induced locomotion relative to WT littermates (p>0.05, Figure S4I), whereas male B6.Del16^+/^*^Bdh1-Tfrc^* mice on the HFD showed significantly attenuated amphetamine-induced locomotion relative to WT littermates (p=0.015, Figure S4J). When the data from STD- and HFD-treated males were directly compared, we found a significant main effect of diet, where males on the HFD showed reduced activity compared to males on the STD (p=0.004), a significant genotype-by-treatment interaction, where male B6.Del16^+/^*^Bdh1-Tfrc^* mice showed reduced activity after amphetamine administration compared to WT littermates (p=0.002), and a significant diet-by-treatment interaction, where males on the HFD showed reduced activity after amphetamine administration compared to males on the STD (p=2.26E-13, Figure S4I-J).

There were no differences in ambulatory activity following saline administration between female B6.Del16^+/^*^Bdh1-Tfrc^* mice on the STD and WT littermates (p>0.05). Likewise, there were no differences in ambulatory activity following saline administration between female B6.Del16^+/^*^Bdh1-Tfrc^* mice on the HFD and WT littermates (p>0.05). After administration of 7.5 mg/kg amphetamine, female B6.Del16^+/^*^Bdh1-Tfrc^* mice on the STD showed significantly attenuated amphetamine-induced locomotion relative to WT littermates (p=9.32E-6, Figure S4K), whereas female B6.Del16^+/^*^Bdh1-Tfrc^* mice on the HFD showed similar levels of amphetamine-induced locomotion relative to WT littermates (p>0.05, Figure S4L). When the data from STD- and HFD-treated females were directly compared, we found a significant main effect of diet, where females on the HFD showed reduced activity compared to males on the STD (p=0.04), a significant genotype-by-treatment interaction, where female B6.Del16^+/^*^Bdh1-Tfrc^* mice showed reduced activity after amphetamine administration compared to WT littermates (p=2.74E-5), a significant diet-by-treatment interaction, where females on the HFD showed reduced activity after amphetamine administration compared to females on the STD (p=6.43E-13, Figure S4K-L).

**Figure S3. Learning phenotypes in STD- and HFD-treated B6.Del16^+/^*^Bdh1-Tfrc^* mice.**

A, B, and C) MWM training A) latency to the hidden platform, B) swim distance, and C) swim speed in STD-treated males (n=5 WT, 8 B6.Del16^+/^*^Bdh1-Tfrc^*).

D, E, and F) MWM training D) latency to the hidden platform, E) swim distance, and F) swim speed in STD-treated females (n=8 WT, 7 B6.Del16^+/^*^Bdh1-Tfrc^*).

G, H, and I) MWM training G) latency to the hidden platform, H) swim distance, and I) swim speed in HFD-treated males (n=12 WT, 12 B6.Del16^+/^*^Bdh1-Tfrc^*).

J, K, and L) MWM training J) latency to the hidden platform, K) swim distance, and L) swim speed in HFD-treated females (n=11 WT, 12 B6.Del16^+/^*^Bdh1-Tfrc^*).

M and N) Percentage of time spent in the quadrant that formerly contained the platform in the probe trial of the MWM in STD- and HFD-treated M) males and N) females.

O, P, and Q) Percent freezing behavior during the fear conditioning O) training phase, P) context test, and Q) tone test in STD-treated males (n=9 WT, 9 B6.Del16^+/^*^Bdh1-Tfrc^*).

R, S, and T) Percent freezing behavior during the fear conditioning R) training phase, S) context test, and T) tone test in STD-treated females (n=9 WT, 9 B6.Del16^+/^*^Bdh1-Tfrc^*).

U, V, and W) Percent freezing behavior during the fear conditioning U) training phase, V) context test, and W) tone test in HFD-treated males (n=12 WT, 13 B6.Del16^+/^*^Bdh1-Tfrc^*).

X, Y, and Z) Percent freezing behavior during the fear conditioning X) training phase, Y) context test, and Z) tone test in STD-treated males (n=12 WT, 13 B6.Del16^+/^*^Bdh1-Tfrc^*).

Data are represented as mean ± SEM. n.s., p>0.05; *, p<0.05; **, p<0.01; ***, p<0.001

Statistical analysis of MWM training phase and fear conditioning (A-L, O-Z) was performed using two-way repeated measures ANOVA. Statistical analysis of MWM probe trial (M and N) was performed using unpaired t-test.

**Figure S4. Acoustic startle, prepulse inhibition, and amphetamine-induced locomotion phenotypes in STD- and HFD-treated B6.Del16^+/^*^Bdh1-Tfrc^* mice.**

A, B, C, and D) Acoustic startle response in A) STD-treated males (n=9 WT, 8 B6.Del16^+/^*^Bdh1-Tfrc^*), B) HFD-treated males (n=12 WT, 13 B6.Del16^+/^*^Bdh1-Tfrc^*), C) STD-treated females (n=10 WT, 9 B6.Del16^+/^*^Bdh1-Tfrc^*), and D) HFD-treated females (n=11 WT, 12 B6.Del16^+/^*^Bdh1-Tfrc^*).

E, F, G, and H) Prepulse inhibition in E) STD-treated males (n=9 WT, 9 B6.Del16^+/^*^Bdh1-Tfrc^*), F) HFD-treated males (n=12 WT, 13 B6.Del16^+/^*^Bdh1-Tfrc^*), G) STD-treated females (n=10 WT, 9 B6.Del16^+/^*^Bdh1-Tfrc^*), and H) HFD-treated females (n=11 WT, 12 B6.Del16^+/^*^Bdh1-Tfrc^*).

I, J, K, and L) Amphetamine-induced locomotor activity in I) STD-treated males (n=9 WT, 9 B6.Del16^+/^*^Bdh1-Tfrc^*), J) HFD-treated males (n=11 WT, 13 B6.Del16^+/^*^Bdh1-Tfrc^*), K) STD-treated females (n=10 WT, 9 B6.Del16^+/^*^Bdh1-Tfrc^*), and L) HFD-treated females (n=9 WT, 12 B6.Del16^+/^*^Bdh1-Tfrc^*).

Data are represented as mean ± SEM. n.s., p>0.05; *, p<0.05; **, p<0.01; ***, p<0.001

Statistical analysis of acoustic startle and amphetamine-induced locomotor activity (A-D, I-L) was performed using linear mixed models. Statistical analysis of prepulse inhibition (E-H) was performed using two-way repeated measures ANOVA.

**Meta-analysis of metabolomics literature**

We performed a meta-analysis of existing literature to test the hypothesis that sex is not commonly addressed as a biological variable in mouse metabolomics studies. A PubMed search for the keywords “mouse metabolomics” yielded 2,601 studies published since 2015. We randomly selected 500 studies for further analysis using a random number generator. We classified the studies based on how sex was addressed as a biological variable; studies were coded as “males only”, “females only”, “sex not specified”, “both sexes, not stratified”, or “stratified”. Of the 500 randomly selected studies, only 44 (8.8%) studied both sexes, and only 17 (3.4%) performed a stratified analysis. 69 studies used only female samples (13.8%), 248 studies used only male samples (49.6%), and 139 studies did not specify the sex of the samples (27.8%).
